# Adolescent environmental enrichment induces social resilience and alters neural gene expression in a selectively bred rodent model with anxious phenotype

**DOI:** 10.1101/2023.10.03.560702

**Authors:** Angela M. O’Connor, Megan H. Hagenauer, Liam Cannon Thew Forrester, Pamela M. Maras, Keiko Arakawa, Elaine K. Hebda-Bauer, Huzefa Khalil, Evelyn R. Richardson, Farizah I. Rob, Yusra Sannah, Stanley J. Watson, Huda Akil

**Affiliations:** Univ. of Michigan, Ann Arbor, MI, USA

**Author notes:** Co-first authorship (Angela M. O’Connor, Megan H. Hagenauer). Corresponding Author: Angela O’Connor.

**Keywords:** environmental enrichment, social stress, adolescence, genetic environment interactions

## Abstract

Stress is a major influence on mental health status; the ways that individuals respond to or copes with stressors determine whether they are negatively affected in the future. Stress responses are established by an interplay between genetics, environment, and life experiences. Psychosocial stress is particularly impactful during adolescence, a critical period for the development of mood disorders. In this study we compared two established, selectively-bred Sprague Dawley rat lines, the “internalizing” bred Low Responder (bLR) line versus the “externalizing” bred High Responder (bHR) line, to investigate how genetic temperament and adolescent environment impact future responses to social interactions and psychosocial stress, and how these determinants of stress response interact. Male bLR and bHR rats were exposed to social and environmental enrichment in adolescence prior to experiencing social defeat and were then assessed for social interaction and anxiety-like behavior. Adolescent enrichment caused rats to display more social interaction, as well as nominally less social avoidance, less submission during defeat, and resilience to the effects of social stress on corticosterone, in a manner that seemed more notable in bLRs. For bHRs, enrichment also caused greater aggression during a neutral social encounter and nominally during defeat, and decreased anxiety-like behavior. To explore the neurobiology underlying the development of social resilience in the anxious phenotype bLRs, RNA-seq was conducted on the hippocampus and nucleus accumbens, two brain regions that mediate stress regulation and social behavior. Gene sets previously associated with stress, social behavior, aggression and exploratory activity were enriched with differential expression in both regions, with a particularly large effect on gene sets that regulate social behaviors. Our findings provide further evidence that adolescent enrichment can serve as an inoculating experience against future stressors. The ability to induce social resilience in a usually anxious line of animals by manipulating their environment has translational implications, as it underscores the feasibility of intervention strategies targeted at genetically vulnerable adolescent populations.

## 1. Introduction

Social stress is a major predictor of future mood disorders, changing both short-term behavioral responses and longer-term developmental trajectories and coping mechanisms [1, 2]. Resilience or susceptibility to social stress shapes how individuals respond to these experiences and the impact on future behavior and brain function [3, 4]. Social stress resilience is determined by an interplay of genetics and environment prior to encountering a stressor [1, 5], and may be modifiable by these same factors [3]. The impact of developmental and environmental factors on resilience and susceptibility to social stress has been studied in mice using behavioral screening [6–9]. Less is known about how genetic predisposition and social temperament influence the experience of social stress, as well as the response to interventions designed to increase social resilience, such as adolescent social experience.

The bred High Responder (bHR) and bred Low Responder (bLR) rat lines robustly model heritable extremes in temperament [10]. Bred based on locomotor reactivity to a novel environment, bHRs exhibit an “externalizing-like” temperament, with disinhibited, hyperactive, and sensation-seeking behavior, while bLRs exhibit an “internalizing-like” temperament, with inhibited, hypoactive, anxious- and depressive-like behavior [10–13]. These bred lines also differ in their response to stressors, including social stress [13–16], enabling insight into the genetic, molecular and circuit mechanisms underlying variable stress responses [13, 17, 18]. Notably, social interaction styles reliably differ between the two lines; bHRs display more aggressive, bold social behaviors and bLRs exhibit more defensive, submissive social behaviors [18, 19]. The extremely stable temperament phenotypes produced by these bred lines enable the study of early-life interventions that may impact lifetime behavioral responses [11].

The divergent temperament in bHR and bLR rats is reflected in differential brain gene expression [20–24], which has persisted across generations and emerges early in life [25]. Recent work uncovered genetic differences that underlie these distinct phenotypes [26, 27]. These brain differences provide insight into behavior beyond our selective breeding paradigm: bHR/bLR differential expression overlaps with that observed in other bred lines targeting similar behavior [24, 25], and predicts the gene expression associated with those behaviors in intercross animals [24] The consistency, stability and predictability of these bred phenotypes across generations [11, 28], along with their innate differences in stress response [13] and social behavior [18, 19] make bLR and bHR animals an excellent model for assessing the interaction of genetics, social stress, and environmental influences.

Environmental enrichment, where animals are housed in complex caging with increased opportunity for sensory stimulation, motor activity and social interaction, can decrease anxiety-like behavior within various animal models [29–33]. Enrichment has long been considered a “eustressor”, the experience of which inoculates against subsequent larger stressors [30, 34]; it is unknown how much genetic vulnerability or stress resilience determines the efficacy of this stress buffering effect. Previous studies in our model indicated that adult environmental enrichment can reduce anxiety-like behavior in bLRs [35] and shift social behavior, decreasing aggression in bHRs and increasing positive-affect ultrasonic vocalizations in bLRs [19].

These previous studies focused on enrichment during adulthood, but adolescence is a critical period for the development of mood disorders, with most mood disorder diagnoses occurring between the ages of 12-18 years [36]. During adolescence, emotional, social, and cognitive circuits undergo a critical period [37–39], allowing social stress reactivity and resilience to be molded by social and environmental conditions [40].

The current study focuses on the impact of adolescent social and environmental enrichment on social interaction, anxiety-like behavior, and social stress resilience in our bred lines to provide insight into how genetics, environment and stress interact. The enrichment conditions used in this study were designed to parse whether the impact of maximal enrichment (social, sensory, and motor) differs from the impact of exposure to social and cage novelty. To characterize the hormonal changes accompanying shifts in social behavior, we measured circulating levels of corticosterone, testosterone, oxytocin, and interleukin-6, all of which have been shown to change with environmental enrichment [19, 41–44], and play a role in mediating social behavior [45–48]. To explore alterations in affective circuitry, we used RNA-seq to measure gene expression within the Nucleus Accumbens (NAcc) and Hippocampus (HC). These brain regions are both impacted by stress [49, 50] and implicated in mediating stress resilience [51, 52] and social behavior [53, 54]. Thus, the current work explored the role of these two important brain regions in mediating the interplay between genetic, developmental and environmental factors in shaping social vulnerability or resilience.

## 2. Methods

Methods are overviewed below, see supplement for details. All animal experiments were carried out in accordance with the National Institutes of Health guide for the care and use of Laboratory animals. All efforts were made to minimize animal suffering.

### 2.1 Experimental Animals

We used male rats from our in-house selectively bred bHR and bLR lines [10] from the generations available at the time of study (F49, F53 and F56). For logistical reasons, sample sizes varied across behavioral tasks and biological measurements (*N*=25-142, *n* per experiment: **Fig S1-S2**). For behavioral outcomes, our study was well powered (80%) to detect medium effects of bred Line, Enrichment, and Defeat, and some of their second-order interactions (d=0.47-0.72) using a traditional alpha (p=0.05), but only sensitive to large (d=0.8-1.2) or very large effects (>1.2) for higher order interactions on behavioral outcomes, and for hormonal and RNA-Seq outcomes.

To decrease litter effects, treatment groups were composed of animals from multiple litters (**Fig S1**). Litters were culled on postnatal day 1 (P1) to even sex ratios (minimum litter size: 3M/3F, maximum: 6M/6F) (**Fig 1A**). Litters were weaned at P21 and males pair- or triple-housed with littermates under a 12:12 hour light:dark cycle (lights on: 7am) with *ad libitum* access to water and food.

**Fig 1:**
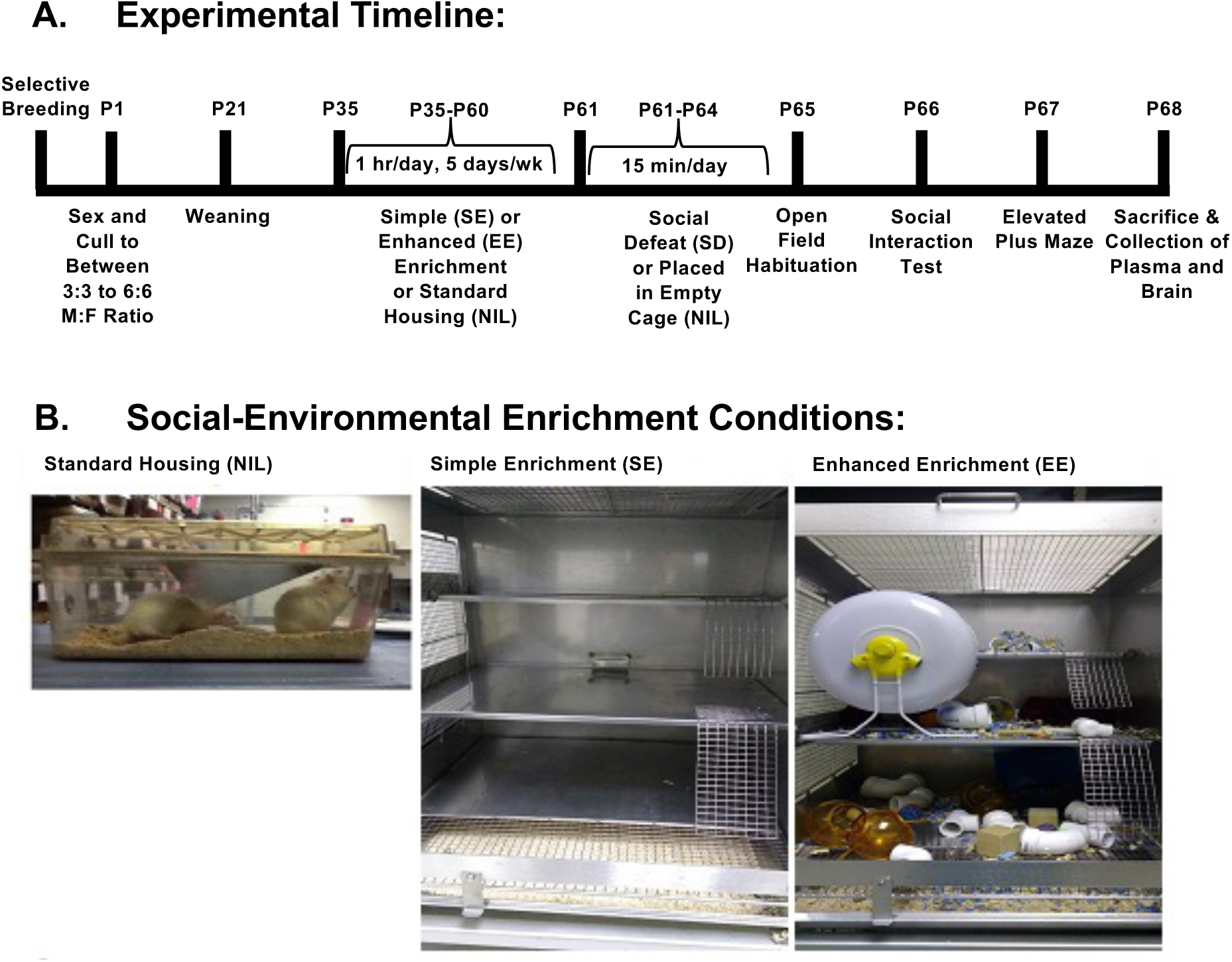
Behavioral Methodology. **A.** Experimental timeline outlining the timing of behavioral interventions and testing. The day of birth is considered postnatal day 0 (P0). **B.** Examples of the standard (“NIL”), simple enrichment (“SE”) and enhanced enrichment (“EE”) cages. Enrichment cages consisted of a large (50×40×50cm) cage with three separate levels connected by mesh ramps. The EE condition also contained various objects that were added to the cage and moved around over the duration of enrichment period, including running wheels, plastic and cardboard tunnels, plastic igloo houses, Nylabones and dog chew toys. Different starting combinations of objects were used each week. All animals from one litter (n=4-6) were placed into the same enrichment cage.

### 2.2 Adolescent Social-Environmental Enrichment

At P35, animals were randomly assigned to standard housing (NIL), simple enrichment (SE) or enhanced enrichment (EE) (**Fig 1A**). SE and EE groups spent an hour a day in large enrichment cages (09:30–10:30 hrs, **Fig 1B**), 5 days/week from P35-P60. All animals from one litter (n=4-6) were placed into the same enrichment cage, mixing siblings from different cages during enrichment. The EE condition also contained various objects (**Fig 1B**) that were added to the cage over each 5-day period and moved daily. All objects were cleaned between weeks (bleach+detergent). The timing and duration of EE followed a protocol akin to [19], with SE included to measure the effects of handling, cage and social novelty.

### 2.3 Repeated Social Stress

The repeated social stress paradigm consisted of a four-day training phase and four-day repeated stressor phase as previously described ([55], see Supplement). All training and social stress took place under red light during the dark period (between 19:00-00:00 hours).

During the training phase, male Long-Evans rats were trained to attack non-experimental outbred Sprague-Dawley intruders, and needed to reach social defeat scores >3 (scores: 1: non-aggressive social interaction; 2: lateral threat and rearing; 3: boxing/scuffling; 4: pinning; 5: pinning and attempted biting) to be used as an aggressor during the stressor phase. Less aggressive Long-Evans (scores <4) served as novel targets in the social interaction test.

From P61-64, the bHR/bLR rats randomly assigned the social defeat group (SD) were introduced individually to a Long-Evans aggressor’s cage for a daily 15-minute stressful social encounter. The bHR/bLR intruder could move freely throughout the cage until an aggressive interaction (score>3). Intruders were then placed into a protective wire mesh container (10×10×15cm) within the Long-Evans’ cage for the remainder of the trial. Each day, each bHR/bLR intruder was exposed to a different aggressor. Any wounding excluded the rat from the study. bHR/bLR rats in the no defeat group (“NIL”) were placed in a clean, empty novel cage within the same testing room and allowed to move freely for the 15-minute period.

Video recordings of behavior during the social stress sessions were hand-scored by a blind observer using The Observer XT software (Noldus Information Technology), with bHR/bLR intruder behaviors classified as submissive or aggressive according to [56, 57], and normalized as a percent of total trial time prior to physical separation from the aggressors (maximum: 15 min).

### 2.4 Social Interaction Test

On P65-66 during the light period (07:00-11:00 hours), bHR/bLR rats underwent social interaction testing consisting of two 5-minute trials on consecutive days (Day 1: habituation, Day 2: testing). The testing arena was a white Plexiglass open field (100×100cm, dim lighting: 40 lux), cleaned (70% ethanol) between animals. bHR/bLR rats were placed into the center of the field. On Day 2, the field included a caged novel stimulus male Long-Evans rat (“target”), with predefined target, interaction, and social avoidance zones. A video tracking system (Ethovision XT 11.5, Noldus Information Technology) calculated the percent time bHR/bLR rats spent in each zone. Precise location and behavior were recorded by a scientist blinded to group status hand-scoring videos (The Observer XT software, Noldus Information Technology).

### 2.5 Elevated Plus Maze (EPM) Test

On P67 during the light period (between 07:00-11:00 hours), bHR/bLR rats underwent the EPM Test. The EPM consists of four intersecting black Plexiglass arms (45cm×12cm) shaped like a cross, elevated 70cm from the floor. Two opposite arms are enclosed (45cm walls) and two remain open; with a square intersection (12×12cm) allowing access to all arms. During the five-minute test, the room was dimly lit (40 lux) and video tracking (Ethovision XT 11.5, Noldus Information Technology) recorded latency to enter the open arms, the amount of time spent in the open arms and centre square. The EPM was cleaned (30% ethanol) between animals.

### 2.6 Tissue and Blood Collection

On P68 during the light period (between 14:00-17:00 hours), bHR/bLR animals habituated to a new room (>30 minutes), then moved to the sacrifice room and immediately decapitated without anaesthesia. Trunk blood was collected in EDTA tubes and placed onto ice before centrifugation (3000 rpm, 4°C, 10 minutes). Plasma supernatant was stored at −80°C. Brains were dissected within 2 minutes of sacrifice, flash-frozen (−30°C), and stored at −80°C.

### 2.7 Corticosterone ELISA

Plasma corticosterone levels were measured using an enzyme immunoassay (EIA) kit (Arbor Assays catalogue# K014-H https://www.arborassays.com/). Following kit protocols, 5μl of plasma from each subject underwent dissociation immediately prior to EIA. Samples and freshly prepared dilution standards were pipetted into well plates in duplicate. A plate reader determined the optical density (450nm) of each well. Corticosterone concentrations were calculated using Arbor Assays’ https://www.myassays.com/. See supplement for other assays (testosterone, oxytocin, interleukin-6).

### 2.8 Behavioral and Hormonal Analysis

All analyses were performed in Rstudio (v.1.0.153, R v. 3.4.1) (code: https://github.com/hagenaue/bHRbLR_Enrichment_Stress_BehaviorAndHormoneData). Non-normal distributions for the dependent variables indicated that inferential statistics were best performed following Log2 data transformation or using non-parametric methods, with batch/generation included as a co-variate. To examine the influence of bred line and adolescent enrichment on behavior during social defeat, we used a full factorial multilevel model that included a linear effect of defeat day (centered on day 4) and ratID as a random effect (function *lme()*, package *nlme* [58], autocorrelation structure: AR1, model fit: maximizing log-likelihood). Results were summarized using *Anova()* (package *car* [59], Type III, contrasts=sum, likelihood ratio test). To examine the influence of bred line, enrichment, and social defeat on other variables, we used a full factorial permutation-based ANOVA (function *aovperm()*, package *permuco* [60], 15,000 permutations, Type III, contrasts=sum). Due to the large number of dependent variables (16), a Bonferonni corrected alpha (adj.p<0.05) defined significance (denoted *). Results at a traditional alpha (p<0.05) are called nominal (denoted #) and considered tentative.

### 2.9 RNA-seq tissue extraction and data processing

All brains were hemi-sected (groups counter-balanced by side), and one half dissected within a cryostat (−20°C). The NAcc (+1.5mm to +1mm Anterior-Posterior [61]) was extracted by 2mm hole-punch, and whole dorsal HC (−3mm to −4mm Anterior-Posterior [61]) extracted using dissection tools.

Nucleotides were extracted from HC and NAcc tissue from a subset of bLR animals (NIL+NIL, NIL+SD, EE+NIL, EE+SD, sample sizes: **Fig S1**), using Qiagen AllPrep DNA RNA miRNA Universal Kit 50. A Nanodrop spectrophotometer quantified the total concentration (285-432 ng/ul) and quality (260/280 ratio: 1.61-1.80) of extracted RNA. Samples were processed by the University of Michigan DNA Sequencing Core (https://brcf.medicine.umich.edu/cores/dna-sequencing/). Samples with RNA integrity numbers (RINs)<8 were excluded (TapeStation automated sample processing system, Agilent, Santa Clara, CA). A DNA library targeting polyadenylated transcripts was constructed for each sample in a 12-cycle PCR (100ng total RNA, KAPA hyper mRNA stranded library prep kit, Roche, catalogue# KK8581). Final cDNA libraries were checked for quality by TapeStation and qPCR (Kapa’s library quantification kit for Illumina Sequencing platforms, catalogue# KK4835, Kapa Biosystems, Wilmington MA). Samples were clustered and sequenced using NovaSeq S2 reagents (NovaSeq S2 Run, Illumina) with 80 samples/flow cell and 50 base paired end reads (targeted sequencing depth=45 million reads/sample).

Reads were aligned to genome assembly Rnor6 (STAR) and summarized into counts per transcript (featureCounts: Ensembl 96 annotation). Following quality control (*see Supplement*), differential expression for the variables of interest (Social Defeat, Enrichment) was calculated using the limma/voom method (*limma* v.3.32.5 [62, 63]) with observed precision weights in a weighted least squares linear regression using models that included region-specific technical co-variates (RNA concentration, RNA extraction batch, dissection batches):

NACC:

*Model 1 (“M1: Main Effects Model”):*

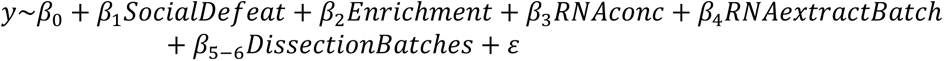

*Model 2 (“M2: Interactive Effects Model”):*

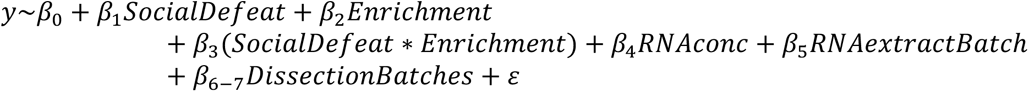

HC:

*Model 1 (“M1: Main Effects Model”):*

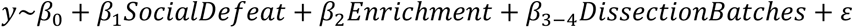

*Model 2 (“M2: Interactive Effects Model”):*

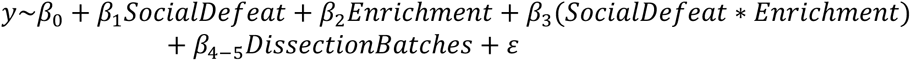

Contrasts were defined by treatment, with the intercept for Enrichment and Social Defeat set as “NIL”. Standard error was moderated using an empirical Bayes distribution (function *eBayes()),* and p-values corrected for false discovery rate (FDR or q-value [64]).

#### 2.9.1 Functional Ontology

We evaluated whether the Enrichment (EE) or Social Defeat (SD) differential expression results (pre-ranked by t-statistic) were enriched for genes representing particular functional, anatomical, and cell-type categories using fGSEA [65] (v.1.2.1, nperm=10000, minSize=10, maxSize=1000) and a custom gene set file (Brain.GMT v.2: 15,545 gene sets). Disproportionate enrichment of significant effects (FDR<0.05 for fGSEA results for M1 or M2 output) within pre-defined categories of gene sets related to our interventions and behaviors was determined using Fisher’s Exact Test. A Bonferonni-corrected alpha (36 comparisons, adj.p<0.05) defined significance. Results at a traditional alpha (p<0.05) are called nominal and considered tentative.

#### 2.9.2 Analysis Code Availability

RNA-Seq data pre-processing was performed using a standard pipeline (MBNI Analysis Hub: https://ahub.mbni.org). All downstream analyses were performed in Rstudio (v.1.0.153, R v. 3.4.1, code: https://github.com/hagenaue/bHRbLR_Enrichment_Stress_RNASeqData).

## 3. Results

We used selectively-bred animals that show an internalizing-like (bLRs) or externalizing-like (bHRs) temperament to examine how genetic background and adolescent social and environmental enrichment interact to shape social behavior, anxiety, and endocrine responses to repeated social stress. We then used RNA-Seq to explore the effects of these interventions on gene expression in the anxious phenotype (bLR) animals in brain regions related to social and emotional behavior and stress regulation (NACC, HC). We discuss the most compelling results below; detailed reporting is in the supplement (**Table S1-S5**).

### 3.1 Behavior during Social Defeat Depends on Genetic Temperament and Adolescent Experience

bHR and bLR rats in the SD group experienced 15 minutes of social defeat stress daily for four days by being placed as intruders into the cage of a larger, territorial Long Evans male. Behavior during these sessions suggested that social defeat stress is not a uniform experience, but instead experienced through the lens of both genetic predisposition and previous social and environmental experience (**Fig 2**). Compared to bHRs, bLRs responded with greater submissive behavior during social defeat (**Fig 2**, Line: X^2^(1, N=70)=88.81, p=2.20e-16*), and bLR submissive behavior increased with each daily defeat session (Day: X^2^(1, N=70)=39.35, p=3.545e-10*; Day*Line: X^2^(1, N=70)=9.91, p=0.00165*) in a manner that was nominally reduced by adolescent exposure to enrichment (Enrichment: X^2^(1, N=70)=8.21, p=0.0165#; Line*Enrichment: (X^2^(1, N=70)=3.42, p=0.0405#). Conversely, bHRs showed more aggressive behavior than bLRs (**Fig 2**; Line: X^2^(1, N=70)=60.99, p=5.735e-15*) in a manner that seemed to potentially increase with each defeat session in animals with previous simple enrichment during adolescence (Day: X^2^(1, N=70)=9.01, p<1e-16*; nominal effects of Day*Line: X^2^(2, N=70)=1.84, p=0.00525#; Enrichment: X^2^(2, N=70)=8.03, p=0.0180#; Day*Line*Enrichment: X^2^(2, N=70)=7.22, p=0.0270#).

**Fig 2.**
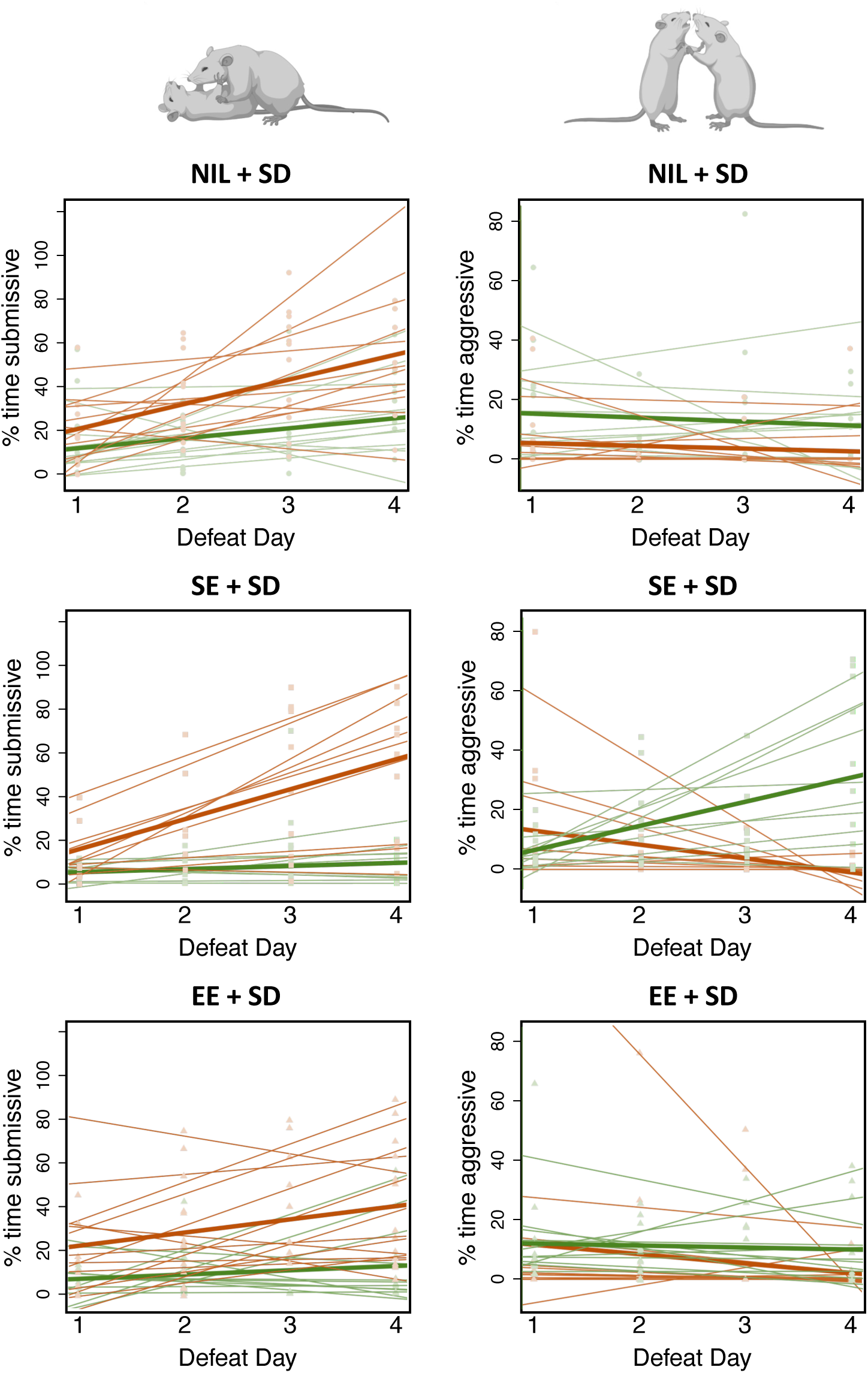
Social Defeat is experienced through the lens of genetic predisposition and potentially previous social and environmental experience. bLR rats responded submissively to social defeat and this increased with each social defeat session in a manner that was nominally moderated by enrichment during adolescence, whereas bHR rats responded aggressively to social defeat in a manner that nominally increased with each social defeat session in animals with previous social experience during adolescence. Bred line is illustrated with color (bHR=green, bLR=red), and adolescent enrichment by datapoint shape (circle=standard housing (NIL), square=simple enrichment (SE), triangle= enhanced enrichment (EE)). Light red or light green dots=individual data points, thin red or green lines: best fit line for each rat across the four days of defeat, thick line: best fit line for the experimental subgroup across the four days of defeat.

Social defeat sessions are complex dyadic interactions, and behavior of resident aggressors may be influenced by intruder phenotype. Indeed, territorial aggressive behavior by the Long-Evans varied depending on whether they encountered a bHR or bLR intruder (**Fig S6**), with the most aggressive intruders receiving more severe defeat (% time aggressive vs. social defeat score: n=242 scored encounters, β=6.305, p=2.00e-16*). Therefore, the more submissive bLRs received lower social defeat scores than the aggressive bHRs (Line: X^2^(1, N=75)=11.17, p=0.000830*, nominal effect of Day: X^2^(1, N=75)=4.65, p=0.0311#). Following full defeat (pinned), a mesh divider was used to prevent injury and further direct interaction. Since aggressive intruders elicited intense territorial aggression, they spent less time directly interacting with the resident Long-Evans (% time aggressive vs. time caged: n=242 scored encounters, β= −0.881, p<2e-16*). Thus, the more-submissive bLRs spent nominally more time caged with their aggressor than the bHRs (Line: X^2^(1, N=71)=5.69, p=0.0170#, Day: X^2^(1, N=71)=8.06, p=0.00453#).

Altogether, we conclude that the experience of social defeat stress differs dramatically for animals that are innately more submissive (bLRs) than animals that are more aggressive (bHRs) and may be modulated by adolescent social-environmental experience.

### 3.2 Exposure to adolescent enrichment altered bLR and bHR social behavior

The impact of our manipulations on social interaction was assessed using a caged novel (neutral) Long-Evans stimulus animal in an open-field arena (**Fig 3A**). Overall, bLRs spent more time in the social avoidance zones than bHRs, away from the stimulus animal (**Fig 3B**; Line: F(1, 128)=22.37, p=6.667e-05*). Previous experience with adolescent enrichment nominally decreased social avoidance in both bHRs and bLRs (**Fig 3B**; Enrichment: F(2, 128)=4.57, p=1.247e-02#).

**Fig 3.**
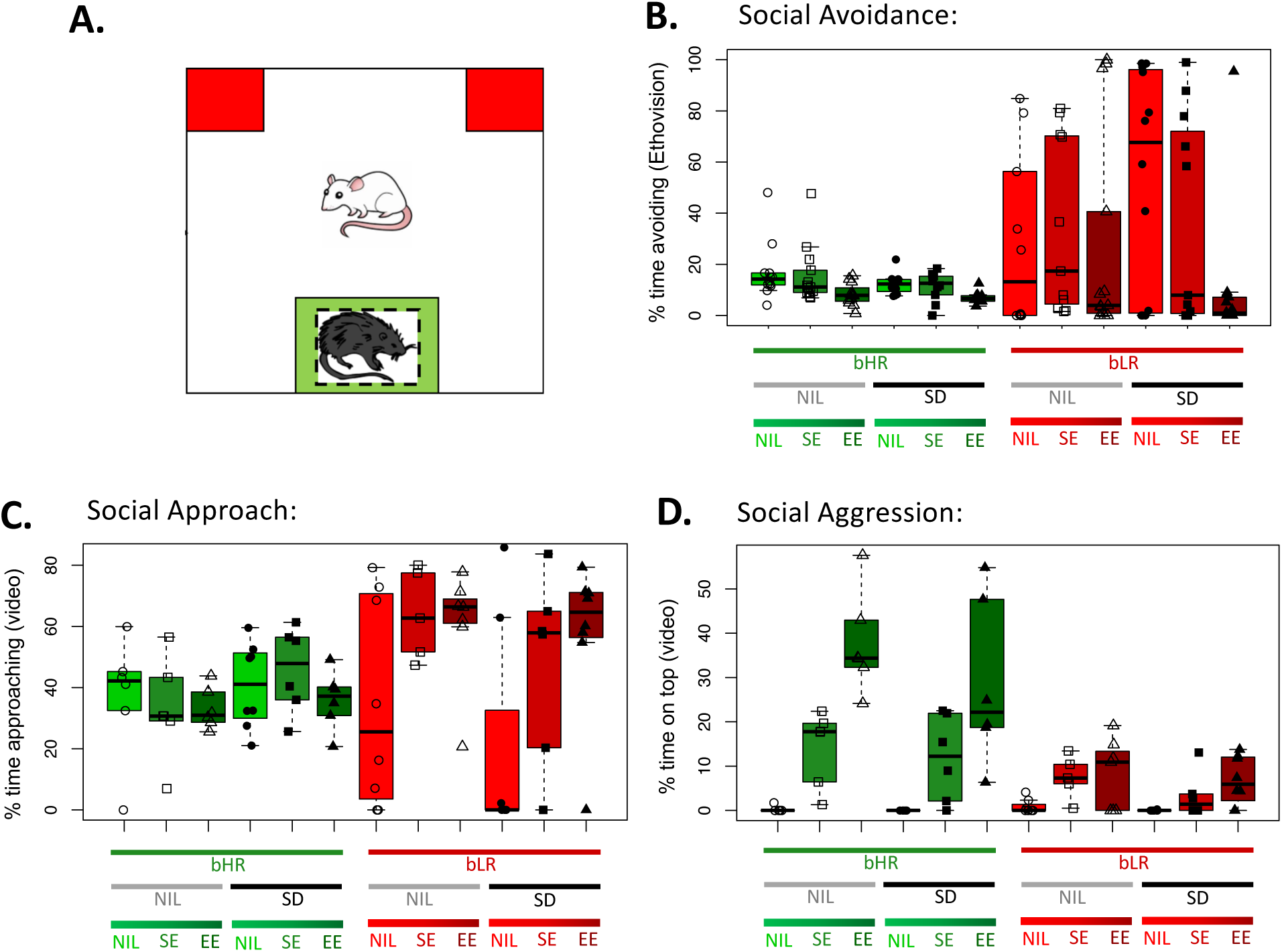
Adolescent social and environmental enrichment seemed to decrease social avoidance, leading to increased approach behavior in bLRs and increased aggression in bHRs. **A.** An illustration of the social interaction task for the bHR/bLR animals (white rat), with each zone delineated: caged novel target Long Evans rat (black rat), avoidance zone (red), and interaction zone (green). **B-D.** Boxplots illustrate the median and interquartile range for each treatment group (+/-whiskers illustrating the range and/or 1.5x interquartile range). Bred line is illustrated with box fill color (bHR=green, bLR=red), adolescent enrichment by datapoint shape (circle=standard housing (NIL), square=simple enrichment (SE), triangle= enhanced enrichment (EE)), and social defeat is indicated by datapoint color (open=no defeat (NIL), black filled=defeated (SD)). **B.** Adolescent enrichment nominally decreased the percent time spent in the socially avoidant zone (as determined by an automated Ethovision analysis). As expected, bLR rats were generally more avoidant than bHR rats. **C.** Adolescent enrichment nominally increased the percent time spent approaching the stimulus animal, especially for bLR rats, as measured by detailed video analysis by a blinded experimenter. **D.** Adolescent enrichment increased the percent time on top of the stimulus animal’s cage, especially for more aggressive bHRs, as measured by detailed video analysis by a blinded experimenter.

Decreased social avoidance did not necessarily translate into pro-social behavior, depending on bred line. Adolescent enrichment increased the percent time spent engaging in social interaction, especially for bLRs, in a manner that seemed to scale with the complexity of the enrichment intervention (NIL<SE<EE, **Fig 3C**). This was nominally evident in a detailed analysis of social approach behavior performed on a subset of the videos (Line*Enrichment: F(2, 67)=4.917, p=0.00940#) and more significantly within an automated Ethovision analysis of time spent in the zone nearby the stimulus animal using videos from all subjects (**Fig S7A**, Enrichment: F(2, 128)=10.51, p=0.0002*).

Adolescent enrichment also increased the percent time spent on top of the stimulus animal’s cage, especially for bHRs, in a manner that scaled with the complexity of the enrichment intervention (NIL<SE<EE, **Fig 3D**). This was evident in a detailed analysis performed on a subset of the videos (Line: F(1, 67)=52.88, p=6.667e-05*; Enrichment: F(2,67)=41.15, p=6.667e-05*; Line*Enrichment: F(2, 67)=21.18, p=6.667e-05*) and within an automated Ethovision analysis quantifying the time spent within the zone on top of the stimulus animal’s cage using videos from all subjects (**Fig S7B,** Line: F(1, 128)=31.26, p=6.667e-05*; Enrichment: F(2,128)=18.14, p=6.667e-05*; Line*Enrichment: F(2, 128)=7.723, p=4.000e-04*). This behavior appeared aggressive, and was often accompanied by loud vocalizations, urinating on the Long-Evans, and biting the bars of the Long-Evans’ cage.

Overall, adolescent enrichment made bLRs more interactive than avoidant, so that they behaved more like typical bHRs, who also displayed increased social aggression following adolescent enrichment. In contrast, social defeat did not have any residual effects on social behavior within this task in either line (p>0.09 for all effects of SD, SD*Line, SD*Enrichment, SD*Enrichment*Line).

### 3.3 Adolescent Enrichment Decreased Anxiety-like Behavior in bHR Rats

On the elevated plus maze (EPM), bHRs and bLRs exhibited expected phenotypical differences [10], with bHRs spending a greater percent time exploring the open arms than bLRs, indicating decreased anxiety-like behavior (**Fig 4A**, Line: F(1, 128)=134.01, p=6.667e-05*). Percent time exploring the open arms was also increased following adolescent exposure to enrichment (Enrichment: F(2, 128)=7.73, p=9.333e-04*), especially in bHRs (Line*Enrichment: F(2,128)=9.668, p=2.000e-04*), in a manner that seemed to scale with the complexity of the enrichment intervention (NIL<SE<EE). Social defeat did not have any residual effects on anxiety-like behavior within this task in either line (p>0.18 for effects of SD, SD*Line, SD*Enrichment, SD*Enrichment*Line).

**Fig 4.**
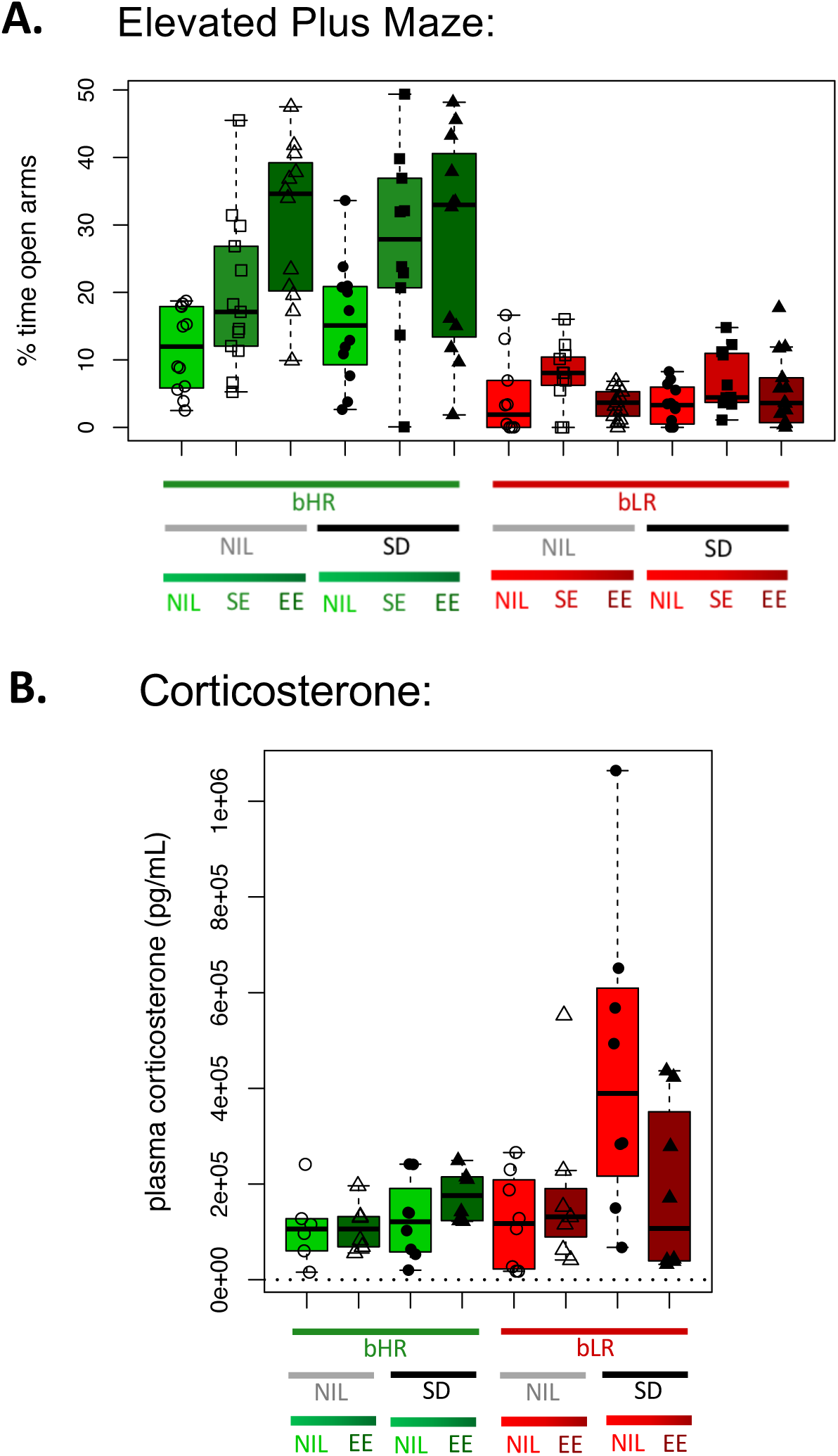
Anxiety and Stress Response: Adolescent enrichment decreased anxiety-like behavior in bHR rats and nominally reduced the elevation of corticosterone following social defeat in vulnerable bLR rats. **A.** The elevated plus maze (EPM) revealed expected phenotypical differences in anxiety and exploratory activity due to selective breeding, with bLRs showing less exploratory activity and elevated anxiety-like behavior, as illustrated by decreased percent time spent in the open arms. bHR rats also showed large increases in the percent time exploring the open arms of the EPM following adolescent enrichment, whereas neither bHR nor bLR rats showed a clear change in anxiety-like behavior following social defeat. Boxplots illustrate the median and interquartile range for each treatment group (+/-whiskers illustrating the range and/or 1.5x interquartile range). Bred line is illustrated with box fill color (bHR=green, bLR=red), adolescent enrichment by datapoint shape (circle=standard housing (NIL), square=simple enrichment (SE), triangle=enhanced enrichment (EE)), and social defeat is indicated by datapoint fill (open=no defeat (NIL), black filled=defeated (SD)). **B.** Plasma corticosterone was nominally elevated at the time of sacrifice several days following social defeat, especially in bLRs kept in standard housing conditions. Note that our hormonal outcome measures were more weakly powered than our behavioral outcomes for logistical reasons: only plasma from a subset of animals was used for this assay (generations F53 and F56: NIL+NIL, NIL+SD, EE+NIL, and EE+SD). The dotted line represents the limit of detection for the assay.

Phenotypical differences in exploratory behavior were also observed when examining the distance travelled during habituation to the open field before social interaction testing (**Fig S8,** Line: F(1,128)=512.26, p=6.667e-05*). There was also a small, nominal decrease in distance travelled following social defeat (Social Defeat: F(2, 128)=6.19, p=1.293e-02#), but no residual effects or interactive effects of enrichment (all p>0.26).

Taken together, these results affirm our selectively-bred model, and provide further evidence that positive effects of adolescent enrichment on behavior depend on genetic predisposition.

### 3.4 bLR Rats Had Nominally Elevated Corticosterone Following Social Stress

To determine whether circulating hormones might contribute to the observed behavioral profiles, trunk blood was collected at sacrifice the day after behavioral testing concluded from a subset of groups representing the greatest range in behavior (generations F53 and F56: NIL+NIL, NIL+SD, EE+NIL, and EE+SD, sample sizes: **Fig S1**). Plasma was assessed for baseline circulating levels of corticosterone, testosterone, oxytocin and IL-6.

Plasma corticosterone was nominally elevated in bLRs compared to bHRs (**Fig 4B**, main effect of Line: F(1, 48)=5.35, *p*=0.0249#). Corticosterone was also nominally elevated in the defeated animals (Social Defeat: F(1, 48)=5.77, *p*=0.0186#), perhaps more so in bLRs kept in standard housing conditions (Line*Social Defeat*Enrichment: F(1, 48)=4.90, *p*=0.0314#). Therefore, exposure to adolescent EE appeared to potentially protect against stress-related corticosterone elevation in bLRs. Interestingly, these nominal effects of social defeat on corticosterone were present in bLRs even though plasma was collected four days after the final defeat session, without evidence of residual social avoidance or anxiety-like behavior.

Testosterone, oxytocin, and IL-6 did not show any convincing effects of treatment group (**Fig S9**-**S10**, *details in supplement*).

### 3.5 Exploratory: Impact of Adolescent Enrichment and Social Defeat on Social-Emotional Circuitry

To better understand the impact of adolescent enhanced enrichment (EE) and social defeat stress (SD) on the more vulnerable bLR rats, we used RNA-Seq to explore gene expression in two brain regions involved in social-emotional processing: the nucleus accumbens (NACC) and hippocampus (HC).

Our NACC and HC RNA-Seq studies were partially independent but revealed surprisingly similar results. Following quality control, the NACC dataset included reads aligned to 17,765 Ensembl-annotated transcripts (median library size=30 million) from 46 subjects (sample sizes: **Fig S1**). Within a model targeting only the main effects of EE and social defeat (M1), five genes were differentially expressed (DEGs) with EE (FDR<0.05), many from the protocadherin family (Pcdhb6, Pcdhga2, Pcdhb5, **Fig 5, Fig S11, Table S2**). There were four DEGs for social defeat (FDR<0.05), many from the Major Histocompatibility Complex (RT1-CE4, RT1-CE5, RT1-N2, **Fig 5, Fig S12, Table S2**). Within a model containing both the main effects and interactive effects of EE and social defeat on gene expression (“M2”), a similar set of DEGs was identified (FDR<0.05: EE: 6 DEGs; SD: 4 DEGs), with no significant interactive effects (EE*SD, all FDR>0.10, **Figs S11-S12, Table S2**).

**Fig 5.**
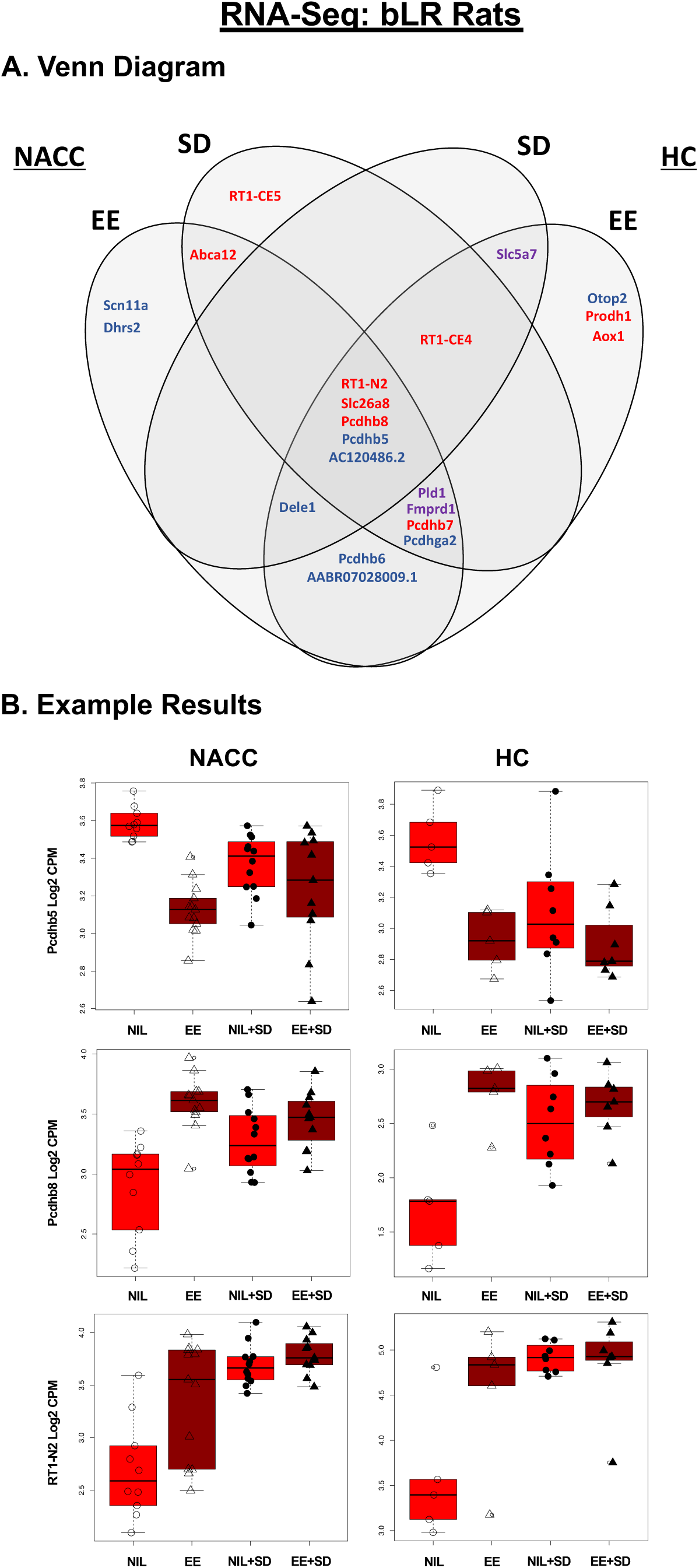
Exploratory bLR RNA-Seq: A similar set of differentially expressed genes (DEGs) were identified in the Nucleus Accumbens (NACC) and Hippocampus (HC) in response to adolescent enhanced enrichment (EE) and social defeat. **A.** A Venn Diagram illustrating the overlap of the bLR RNA-Seq results from the two brain regions (NACC, HC) and treatment groups (adolescent enrichment: standard housing (NIL) vs. enhanced enrichment (EE); social defeat: no defeat (NIL) vs. social defeat (SD)). To be included in the Venn Diagram, a gene needed to be differentially expressed in association with either EE or SD in at least one region (FDR<0.05). Then, to be considered overlapping, there needed to be at least nominal (p<0.05) differential expression with the other intervention in the same brain region, or in association with either intervention in the other brain region. Surprisingly, the overlapping effects of EE and SD were often in the same direction in both brain regions: Red=upregulation, Blue=down-regulation, Purple=differential expression in opposing directions under different conditions/regions. For the full table of top DEGs (FDR<0.05) see **Figs S12-14**. For the full results for all genes see **Tables S2-S3**. B. Example boxplots illustrating the relationship between gene expression (log2 CPM) and treatment group. Adolescent enrichment is illustrated by datapoint shape (circle=standard housing (NIL), triangle=enhanced enrichment (EE)) and social defeat is indicated by datapoint fill (open=no defeat (NIL), black filled=defeated (SD)). Please note that these results should be considered exploratory, as low statistical power can disproportionately increase false positive risk when using discovery-based approaches.

The HC dataset was smaller and had a shallower read depth. Following quality control, the HC dataset had reads aligned to 17,629 Ensembl-annotated transcripts (median library size=20 million) from 25 subjects (sample sizes: **Fig S1**). Within both models (M1 and M2), there was a similar set of DEGs for EE (FDR<0.05, M1: 6 DEGs, M2: 1 DEG, **Fig S13, Table S3**) and no significant DEGs for social defeat or interactive effects of social defeat and EE (all FDR>0.10**, Table S3**).

Notably, there was substantial overlap between the DEGS that were identified in response to the two interventions within both brain regions (**Fig 5).** Most DEGs for either EE or social defeat in one region (FDR<0.05) showed at least nominal effects (p<0.05) with the other intervention in the same brain region, or with either intervention in the other brain region. These overlapping effects of EE and social defeat were often surprisingly in the same direction in both brain regions (**Fig 5, Figs S11-S13).**

To explore the brain functions associated with this differential expression, we compared our results to a custom gene set file (Brain.GMT v.2) that included not only traditional gene ontology, but also gene sets from previous publications and public data releases related to brain cell type, co-expression networks, stress, behavior, and neuropsychiatric illness. Using gene set enrichment analysis, we found that differential expression was enriched within a disproportionately large percent of the gene sets in categories paralleling our behavioral results: stress, fear conditioning, social behavior, aggression, and activity level (**Fig 6**, full results: **Table S4-S5**). These enriched gene sets included many of our top DEGs for adolescent EE (*Scn11a, Pchdb5, Pchdb8, Prodh1*) and social defeat (*RT1-CE4, RT1-CE5, Abca12, Slc5a7*)(**Figs S12-14**). Given this converging evidence, these DEGs might be particularly worthy of future study in association with behavior. Within these gene sets, adolescent EE and social defeat typically had opposing effects within the NACC and similar effects within the HC (**Fig S14-19).** This is interesting, because although environmental enrichment and social defeat are often considered opposing interventions in terms of their effects on stress susceptibility, both interventions involve a novel environment, social stimuli, increased activity, and some amount of stress.

**Fig 6.**
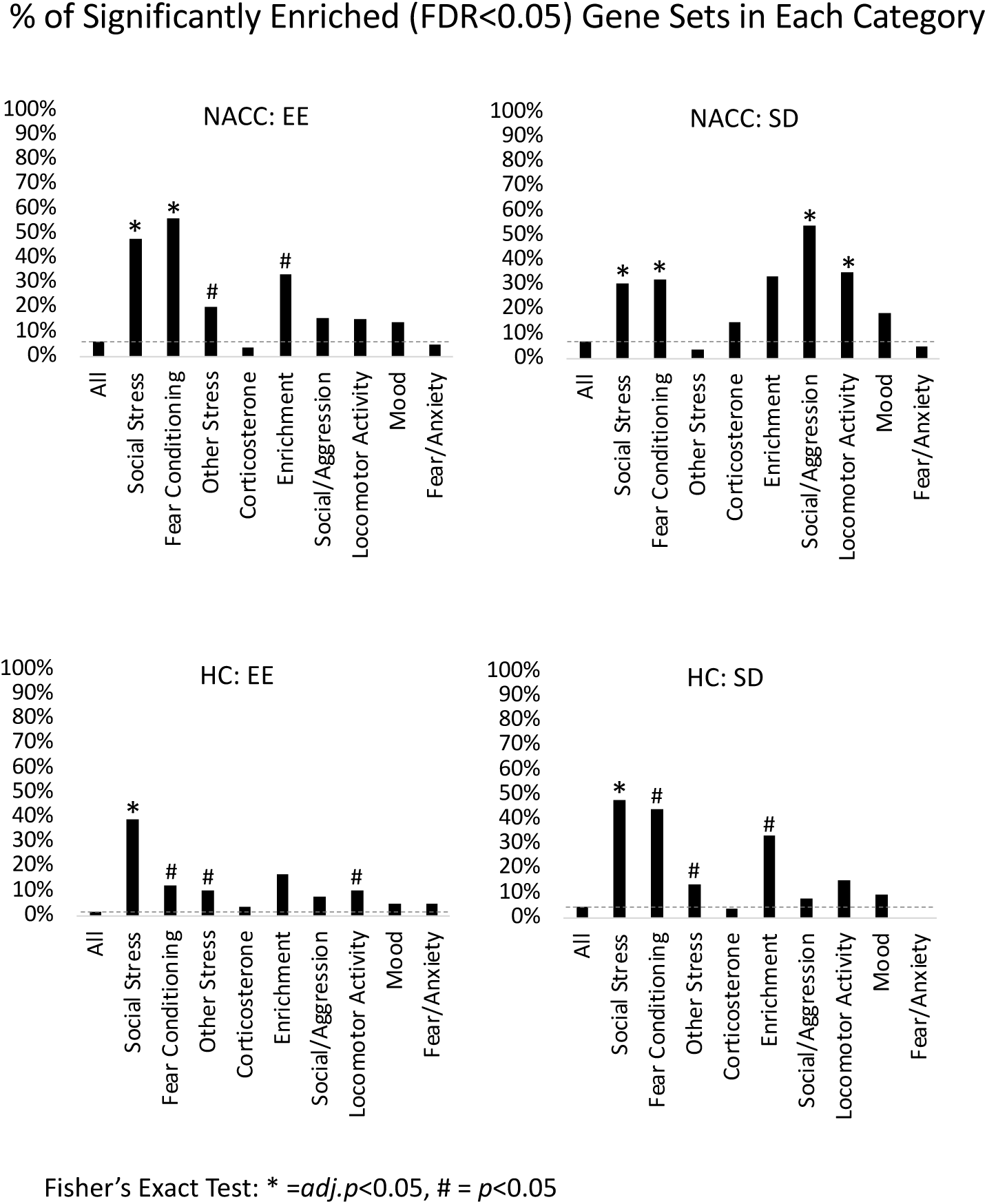
Exploratory: For each of our interventions, differential expression was enriched within gene sets related to stress, social behavior, aggression, activity level, and potentially environmental enrichment. Using gene set enrichment analysis (fGSEA) and a custom gene set file (Brain.gmt), we found that there was enriched differential expression within a disproportionate percent of the gene sets in categories paralleling our behavioral results. Shown above are the percent of gene sets (y-axis) from each category (x-axis) that were found to be significantly enriched with differential expression (FDR<0.05 using fGSEA output from the main effects model (M1) or interactive effects Model (M2)) for each intervention (enhanced enrichment (EE), social defeat (SD)) in each brain region (nucleus accumbens (NACC), hippocampus (HC)) (subpanels). Categories for which a disproportionately large percent of gene sets were enriched with differential expression were identified using fisher’s exact test: *adj.p<0.05, #p<0.05. The directionality for individual enriched gene sets can be seen in **Figs S14-S19**. For the full results for all gene sets, see **Tables S4-S5**.

Differential expression was also enriched in many gene sets related to cell type (**Figs S20-S21**). Gene sets related to oligodendrocytes were enriched with upregulation in both the NACC and HC following both social defeat and adolescent EE, perhaps implying increased connectivity. Other cell type related gene sets showed divergence across conditions: in the NACC, neuronal gene sets were enriched with down-regulation following social defeat, and gene sets related to brain vascular and ventricular systems (astrocytes, mural cells, endothelial cells, progenitor cells, neurogenesis-related cells, ependymal cells) were enriched with down-regulation following EE. In the HC, choroid plexus gene sets were upregulated following social defeat and EE.

In conclusion, we found that adolescent EE and social defeat may impact social and emotional neural circuitry and associated genes in a manner that is likely to reflect features shared across these social-behavioral interventions, and may be accompanied by structural as well as functional changes.

## 4. Discussion

The present results extend our understanding of how genetic predisposition influences individual responses to interventions intended to increase social resilience, on a behavioral, hormonal, and neural level. This is the first study to examine adolescent enrichment in combination with repeated social stress in a model with heritable differences in temperament. The bLR and bHR rodent lines are a well-characterized, stable model of genetic contributions to behavioral temperament [11, 66–69][25, 26], which makes them ideal subjects for examining early-life interventions. We found that a month of adolescent social and environmental enrichment produced long-term effects on social behavior, aggression, and anxiety, that varied according to genetic background and temperament. In bHRs, adolescent enrichment increased aggressive behavior in response to neutral social encounters and possibly territorial aggression. In bLRs, enrichment appeared to decrease submission in response to territorial aggression and increased social approach during a neutral social encounter. Physiologically, stress hormone appeared elevated in standard-housed bLRs that had experienced social defeat, which seemed reduced in bLRs that had adolescent enrichment, suggesting a lasting buffering effect of adolescent intervention. Therefore, following adolescent enrichment, the typically anxious bLRs appeared to develop greater resilience to social defeat and their social interactions came to resemble behavior more typical of bHRs. Thus, adolescent social and environmental complexity may be a promising means for inducing social resilience in vulnerable individuals.

Focusing on the anxious bLRs, we conducted an exploratory RNA-Seq study quantifying the impact of adolescent enrichment and social defeat stress on gene expression in two brain regions involved in affect, stress regulation, social behavior, emotional memory, and behavioral inhibition: the NACC and HC. Differential expression was disproportionately enriched within gene sets related to stress, social behavior, aggression, and exploratory activity [49–54]. Additionally, adolescent enrichment and social defeat often had similar effects on gene expression, emphasizing that although these interventions are considered to have opposite effects on stress susceptibility, they share many similarities, including novelty, social interaction, exploration, and stress.

### 4.1 Behavior and Hormones

In mice, adolescent enrichment preceding social defeat stress primarily impacts social behavioral outcomes [70]. Similarly, adolescent enrichment increased social interaction in outbred Sprague-Dawley rats [71] and adult enrichment altered social behaviors in bLR/bHR rats, increasing positive-affect ultrasonic vocalizations and decreasing bHR aggressive behaviors [19]. bHRs typically show greater social interaction than bLRs [67]; our current findings show that adolescent enrichment may increase bLR social approach and interaction. This could be due to motivation to exert social dominance over a restrained Long-Evans target, as both bLRs and bHRs spent more time on top of the stimulus animal’s cage during social interaction testing.

In mice, social enrichment alone has less impact than social-environmental enrichment [75]. In our current study, simpler enrichment (SE) produced fewer changes in bLR social behavior compared to enhanced enrichment conditions (EE), suggesting that the extra physical experiences available in the EE cage might be important for producing long-term changes in social behavior in our usually anxious line. Both types of enrichment increased motor activity due to the larger cage size, but EE included a running wheel and a variety of toys/objects. Increased motor activity alone is known to impact hippocampal neurogenesis and animal behavior [76, 77]. Combined environmental enrichment (social, sensory and motor) was used in this work as it is thought to be maximally stimulating and rewarding to animals [76, 78].

In Sprague-Dawley outbred rats, adult enrichment increases stress resilience [79], and adolescent enrichment alters the neural circuitry underlying stress responsiveness [71]. In mice, the impact of enrichment on resilience to social stressors depends on genetic background and baseline aggression [72–74]. Adult enrichment also decreased anxiety-like behaviors in outbred Sprague-Dawleys [31], Roman Low Avoidance rats [33], bLR and bHR rats [35]. Our current results confirm that adolescent enrichment decreases bHR anxiety-like behavior, but did not appear to alter bLR anxiety-like behavior within our experimental timeline.

We also did not observe the typical increase in social avoidance and anxiety behaviors seen following social defeat [84]. For bLRs, effects may have been obscured by their already profoundly anxious and submissive behavior hitting a floor on behavioral assays. However, the lack of effect of social defeat on bHR behavior or bLR behavior during the social interaction test is not as easily dismissed, especially since our sample for these outcomes was well-powered, and suggests that our social defeat intervention should be considered mild, either due to the lack of 24 hr exposure to the aggressor as is typical in some paradigms [70] or due to protective factors in our design, such as social housing.

Despite the minor impact of social defeat on measured behavioral outcomes, there appeared to be systemic effects of social stress in bLRs that were reduced by adolescent enrichment. Corticosterone was nominally elevated in socially defeated bLRs at sacrifice, which occurred at a time close to the daily nadir in corticosterone release, when corticosterone levels are sensitive to both depression and stress experiences [80–82]. As sacrifice occurred four days after social defeat, these results could suggest persistent elevation, or perhaps increased reactivity to subsequent testing and handling following social stress. This stress-related elevation in corticosterone appeared reduced in bLRs following adolescent enrichment. Our results parallel previous findings that repeated adolescent social stress increases corticosterone in outbred Sprague-Dawley rats [83], but contrast with the increased corticosterone observed in bHRs immediately after extended exposure to adult enrichment [19] and the enhanced stress response observed in outbred rats after adult enrichment [79]. Altogether, these findings suggest that exposure to enrichment can act as an eustressor [30, 34, 79], increasing stress responses acutely but decreasing them in the long-term.

These findings are important because inducing resilience in the anxious bLRs has proven difficult, usually requiring pharmacological intervention [28, 35]. bLRs that experienced adolescent enrichment showed increased social interaction, and nominally decreased submission during social defeat, decreased social avoidance, and a reduced elevation in corticosterone following social stress. This increased social resilience developed despite the relatively short exposure to enrichment in our paradigm (1 hr/day vs. a typical 24 hrs/day), suggesting that adolescence may be a potent time for intervention or that the daily removal of enrichment may enhance its benefits as a eustressor. Other studies have similarly observed large changes in baseline bLR behaviors following adolescent experiences [85, 86]. Recently, adolescent enrichment preceding repeated social stress was investigated in mice, showing similar positive effects [70]. In contrast, removal of adolescent enrichment elicited a depression-like phenotype in outbred Sprague-Dawley rats [78]; however, these animals were single-housed for two weeks following loss of enrichment, a housing condition that alone induces stress responses [87, 88]. Our bLR and bHR animals were never single-housed. Altogether, the large impacts of enrichment exposure and removal during this life period suggest that adolescence is a time of both great vulnerability and potential for positive intervention.

### 4.2 RNA-seq

RNA-seq was used to explore how adolescent enrichment and social defeat might impact the NACC and HC, two regions important for stress and social behavior [49–54]. We observed differential expression related to both interventions, despite the minor impact of social defeat on measured behavioral outcomes. Previous work using a subset of samples taken from our study also found an increase in a pro-depression growth factor in defeated bLRs [89]. Notably, our findings suggest that the most impactful aspects of our two interventions on brain function may not be their assumed affective valence (positive vs. negative) but shared characteristics, such as repetitive exposure to social stimuli and a complex environment. The mild nature of our defeat paradigm and timing of brain collection may also have contributed to a similar impact of adolescent enrichment and social defeat stress on gene expression. Previous work demonstrated that removing enrichment for a week elicits a stress response [78, 90], which is similar timing to when we collected brain tissue, although we did not observe elevated corticosterone in our enriched groups.

Our exploratory RNA-Seq analysis provides promising molecular targets for future studies. Two groups of genes were particularly impacted by both interventions in both brain regions: the RT1 and protocadherin genes. The RT1 genes are part of the class III region within the rat major histocompatibility complex, important for producing cytokines and complement components [91]. Changes in cytokine expression regulate both social behavior and neuronal connectivity [92], while social status and experience impact cytokine levels in multiple species [93–96]. Living in a social group is an immune challenge, and there is strong cross-talk between immune and social signalling pathways [97, 98]. The current data provide further evidence linking immune and social signalling and suggest that both positive and negative social experiences regulate immune signalling within the brain.

Protocadherins are part of the Cadherin family of transmembrane glycoproteins that regulate cell-to-cell contact through extracellular interactions [99]. We observed differential expression within the class of clustered protocadherins [100], which regulate neurite formation, including dendritic self-avoidance, arborization, spine formation, axonal branching, and pruning [101–104]. Many of these protocadherins were down-regulated in both brain regions following adolescent enrichment and sometimes social defeat. Previous research found that adult enrichment decreased other brain glycoproteins [105, 106] that regulate neuroplasticity [107–109], reopening critical periods within the brain [106]. Protocadherins also regulate neuroplasticity and development [110, 111]; thus, adolescent enrichment and social defeat might similarly extend or reopen plasticity gated by protocadherin expression.

Using gene set enrichment analysis, we found that differential expression was disproportionately enriched in categories paralleling our behavioral results: gene sets related to social behavior, aggression, and activity level. Therefore, our current study not only provides evidence that adolescent enrichment and social stress impact social behavior, but also implicates gene expression networks that mediate these behaviors. Differential expression was also enriched within gene sets associated with fear conditioning and other stressors. While intuitive, these findings may be due to the prevalence of immediate early genes within these gene sets rather than a “stress network” specifically.

Both adolescent enrichment and social defeat increased expression within oligodendrocyte-related gene sets within both brain regions. Previous studies found that enrichment increased myelination and oligodendrocyte markers and expression in mice, particularly during adolescence [112, 113]. Our results suggest that this effect also occurs in rats. The impact of social stress on oligodendrocytes and brain myelination is less clear-cut, and may depend on brain region, stress susceptibility [114–117], and potentially species. In male adolescent Sprague Dawley rats, increased myelination within the hippocampus was similarly observed following juvenile stress [118].

There were opposing effects of adolescent enrichment and social defeat stress on the expression of ventricular and endothelial-related gene sets within the NACC. In mice, social stress similarly increased brain endothelial and ependymal markers [119–121], suggesting an increased inflammatory state. The effects of adolescent enrichment on these supporting cells and brain systems are less straightforward, varying with the timing and duration of enrichment, species/strain, and brain region [122–125]. We observed decreased expression of ventricular and endothelial-related genes within the NACC following adolescent enrichment in bLR rats.

Altogether, the gene expression profiling results underscore the multiple classes of mechanisms that could participate in resilience induction in the bLRs, including cell type balance, immune signalling, and neuroplasticity. Although our findings should be considered exploratory until replicated, the overlap in affected genes and gene sets between both brain regions provides reassurance that our differential expression models properly controlled for dataset-specific confounding technical variability. Similarly, many of our top DEGs were in gene sets previously implicated in stress and behavior (**Figs S12-14**), and our findings add to a growing body of evidence implicating immune signalling, glycoproteins, myelination, and vascular-related cell types in enrichment-related neuroplasticity and social behavior. Although we did not include bHRs in our RNA-Seq analysis, previous findings suggest that some of these pathways may differ in our bred lines (**Fig S22**), indicating that it is worthwhile to design studies examining the impact of interventions in animals with different genetic vulnerabilities.

### 4.3 Limitations and Future Directions

One notable limitation of our study is that it only used males, as the social defeat paradigm is best characterised in male rodents; thus any interactions between the effects of environmental enrichment and social stress in females may involve both shared and distinct mechanisms. Another limitation to our study is the possibility that some results may be partially driven or obfuscated by litter effects, which are known to particularly influence male adolescent rat social behaviour [126]. Our design relied on the inclusion of siblings in experimental groups, but we included rats from multiple litters and multiple generations in each of group to minimize effects on behavioral profiles.

### 4.4 Conclusion

Repeated social stress is often used to model depression and anxiety; our findings indicate that social defeat is not a uniform experience but should instead be considered through the lens of genetic predisposition and previous social and environmental experience. Our findings support the concept that enrichment serves as a “eustressor”, providing a mild inoculating dose of stress due to novelty that improves future coping with larger stressors. We find that adolescent enrichment influences future social interactions and anxiety-like behavior in a manner that depends on genetic temperament. Exploratory RNA sequencing in the HC and NACC in vulnerable bLRs further suggests that social-emotional circuitry and associated gene families are altered following adolescent enrichment and social defeat, encoding both the similar and divergent aspects of these social-behavioral interventions. The ability to potentially induce social resilience in a usually anxious line of animals by manipulating the adolescent environment provides an exciting avenue for the development of interventions targeted at vulnerable human adolescent populations.

## Supporting information

Supplemental Methods and Results

## Credit authorship contribution statement

**Angela M. O’Connor:** Conceptualization; Data curation; Formal analysis; Investigation; Methodology; Project administration; Validation; Visualization; Roles/Writing - original draft; Writing - review & editing. **Megan H. Hagenauer:** Conceptualization; Data curation; Formal analysis; Investigation; Methodology; Project administration; Resources; Software; Validation; Visualization; Roles/Writing - original draft; Writing - review & editing. **Liam Cannon Thew Forrester:** Data curation; Formal analysis; Methodology; Software; Writing - review & editing. **Pamela M. Maras:** Investigation; Methodology; Writing - review & editing. **Keiko Arakawa:** Investigation; Methodology; Writing - review & editing. **Huzefa Khalil:** Data curation; Formal analysis; Software; Writing - review & editing. **Evelyn R. Richardson:** Data curation; Formal analysis; Software; Writing - review & editing. **Elaine K. Hebda-Bauer:** Resources; Writing - review & editing. **Farizah I. Rob:** Data curation; Formal analysis; Software; Writing - review & editing. **Yusra Sannah:** Data curation; Formal analysis; Software; Writing - review & editing. **Stanley J. Watson, Jr.:** Conceptualization; Funding acquisition; Project administration; Resources; Supervision; Writing - review & editing. **Huda Akil:** Conceptualization; Funding acquisition; Project administration; Resources; Supervision; Writing - review & editing.

## Acknowledgements

We gratefully acknowledge the help of James Stewart and Alexander Stefanov during animal tissue collection, and Fei Li for assistance with animal behavior.

## Data Availability

All data are released on Figshare (DOI: 10.6084/m9.figshare.24085524) and GEO/SRA (Accession# GSE237890, https://www.ncbi.nlm.nih.gov/geo/query/acc.cgi?acc=GSE237890).

