## Supplemental Methods and Results for "Adolescent environmental enrichment induces social resilience and alters neural gene expression in a selectively bred rodent model with anxious phenotype"

### Table of Supplementary Figures

Fig S 1. The subgroup sample sizes associated with each dependent variable. 9
Fig S 2. The subgroup sample sizes with video footage for each day of social defeat. 10
Fig S 3. Histograms showing that most of the dependent variables had a non-normal distribution. 11
Fig S 4. RNA-Seq quality control: Library size varies with treatment group. 15
Fig S 5. RNA-Seq quality control: Identifying important technical covariates for the differential expression model using principal components analysis. 16
Fig S 6. The resident Long-Evans aggressor animals appeared to respond to aggressive behavior during social defeat sessions with more severe defeat (social defeat scores>4), leading to a nominally earlier intervention by the experimenter to stop direct interaction (time caged lower than the full 15 minutes). 20
Fig S 7. Adolescent social and environmental enrichment increased approach behavior in bLRs and increased aggression in bHRs: automated Ethovision analysis. 21
Fig S 8. Exploratory behavior in an open field revealed expected phenotypical differences due to selective breeding, but potentially unexpected effects of social defeat and adolescent enrichment. 23
Fig S 9. Plasma testosterone is elevated in bLR rats, potentially more so when kept in standard housing conditions. 24
Fig S 10. There were no significant effects of treatment group on basal plasma hormone levels of oxytocin or IL-6. 25
Fig S 11. The top genes differentially expressed in relationship to EE in the NACC (FDR<0.05) often show a similar pattern of effects in the HC. 27
Fig S 12. The top genes differentially expressed in relationship to Social Defeat (SD) in the NACC (FDR<0.05) often show a similar pattern of effects in the HC. 29
Fig S 13. The top genes differentially expressed in relationship to Enhanced Enrichment (EE) in the HC (FDR<0.05) often show a similar pattern of effects in the NACC. 31
Fig S 14. Gene sets that were previously associated with social behavior, aggression, and locomotor activity are enriched with down-regulated expression related to Social Defeat (SD) in the NACC. 32
Fig S 15. Gene sets that were previously associated with increased locomotor activity are enriched with upregulation related to Enhanced Enrichment (EE) and Social Defeat (SD) in the HC, and gene sets associated with decreased locomotor activity are enriched with down-regulation. 33
Fig S 16. Gene sets that were previously associated with social stress, fear conditioning, and other stress are highly enriched with differential expression related to Enhanced Enrichment (EE) and Social Defeat (SD) in the NACC. 34
Fig S 17. Gene sets that were previously associated with social stress, fear conditioning, and other stress are highly enriched with differential expression related to Enhanced Enrichment (EE) and Social Defeat (SD) in the HC. 35
Fig S 18. A few gene sets related to mood/depression and endogenous fear & anxiety were enriched for differential expression related to Social Defeat (SD) and Enhanced Enrichment (EE) in the NACC. 36
Fig S 19. A few gene sets related to mood/depression and endogenous fear & anxiety were enriched for differential expression related to Social Defeat (SD) and Enhanced Enrichment (EE) in the HC. 37
Fig S 20. Cell type related gene sets are enriched with differential expression in the NACC associated with SD and EE. 38
Fig S 21. Cell type related gene sets are enriched with differential expression in the HC associated with SD and EE. 39
Fig S 22. Some of the top differentially expressed genes in our study have been previously shown to have nominal (p<0.05) bLR/bHR differential expression in the hippocampus. 40

### Table of Supplementary Tables

### Supplementary Methods

#### bHR and bLR breeding protocols

The bred High Responder (bHR) and bred Low Responder (bLR) lines are well-established rodent models of divergent temperament, bred based on reactivity to a novel environment [1]. Animals are bred on site in the research facility; both selectively bred lines are maintained in 12 distinct "family" groups, with rotational breeding within each of the two lines to minimize in-breeding, as is best practice when maintaining a selectively bred animal line [2]. All colony animals are tested for locomotor activity between P55-P65, and bred pairs are composed of males and females with similar locomotor activity scores. A proportion of animals in each generation are used to maintain the two colonies and family lines; researchers are allocated breeding pairs on request, and the offspring used in experimental procedures. Males and females are paired for 10 days, and females monitored for signs of pregnancy. Animals with locomotor activity scores that diverge too far from the average for their bred line are not used in either colony breeding or experimental pair breeding.

#### Animal Housing and Husbandry

Upon weaning, experimental animals were housed on a 12:12 hour light:dark cycle (lights on: 7am, light intensity 50-100 Lux depending on row level within cage racks) at 21-25$^{o}$C and 30-50% humidity with *ad libitum* access to water and food (5LOD, PicoLab Laboratory Rodent Diet, LabDiet pellets). Experimental bLR/bHR animals were never housed in the top row of the caging racks; experimental groups were randomly distributed throughout the caging rack and always returned to the same shelf position following any testing. Following the completion of adolescent enrichment and prior to beginning social defeat stress, all experimental animals were marked on their tails with identifying numbers to allow researchers to track the behavior of individual animals, which was the experimental unit used in our study. Cages were kept in an animal biosafety level-1 housing room, and changed once a week, with water bottles changed twice a week. Cage bedding consisted of corn cob pellets (bed-o'cobs, The Andersons, Lab Bedding Products), and animals were housed in pairs or triples with litter mates. All bLR/bHR experimental animals were housed in the same room for all three generations tested. The Long-Evans rats used as aggressors for the social defeat task were always housed in a separate room to experimental animals, which was the same room for housing all Long-Evans rats.

#### Experimental Procedures: Transport, Acclimation, Timing, Order, and Experimenters

Cages were transported between housing rooms and behavioral testing rooms using a cart. During social defeat stress, Long-Evans aggressors were placed in the behavioral room for half an hour before beginning social defeat, to give them time to settle and adjust in a new environment to encourage territorial behavior. bLR/bHR experimental animals remained within their housing room until immediately before undergoing social defeat, and were placed straight back into their housing room after finishing their defeat session, to prevent prolonged exposure to the Long-Evans aggressors and also prevent them from overhearing other animals experiencing social defeat and developing extra stress as a result. During behavioral testing, bLR/bHR experimental animals were placed into the behavioral testing room half an hour before experiments started, to give them time to settle after being moved and prevent testing order effects. The order that the rats underwent each behavioral test was randomly selected (i.e., which rat was tested first, which rat was tested second…), with animals run in different orders on each day of testing.

Social defeat stress was conducted at night, during the animals' awake period, to maximize territorial aggressive behavior from the Long-Evans aggressor rats; behavioral testing was carried out on the bLR/bHR animals in the morning as per established laboratory protocol. At sacrifice, bLR/bHR experimental animals were placed into a holding room next door to the sacrifice room for half an hour after being moved from their housing room, to decrease the stress effects of being moved through the animal house. The order that the rats were sacrificed was assigned randomly, to prevent potential stress impacts on any particular experimental group. Individual animals were moved by hand from the holding room into the sacrifice room, where they were immediately decapitated.

The same female experimenter carried out all social defeat stress and behavioral testing, wore the same set of provided scrub clothes during live animal experiments, and maintained the same fragrance profile (laundry detergent, deodorant, shampoo, body wash and body lotion) across all animal generations tested. The experimenter had no pets at the time of these experiments, and was assisted by the same two male research technicians during all sacrifices.

#### Repeated Social Stress

##### Resident/Aggressor Animals

All training and social stress took place under red light during the dark phase of the light cycle (between 19:00-00:00 hours). During both phases, Long-Evans male rats (400–500g) were housed with ovariectomized female rats to increase their territorial response and aggressive behavior. Experimental bHR and bLR animals and Long-Evans resident aggressors were housed in separate rooms throughout the entirety of testing. Adult, retired-breeder Long-Evans male and female animals were purchased from Taconic and Charles River. Females underwent ovariectomy surgery followed by a recovery period of 14 days. Male and female Long-Evans rats were then pair-housed for two weeks prior to training and the repeated social stressor. During the training phase, male Long-Evans were trained to attack non-experimental outbred Sprague-Dawley intruders and were used as aggressors during social defeat stress if aggression scores of at least 4 were reached (aggression scores: 1 non-aggressive social interaction; 2 display of lateral threat and rearing; 3 boxing/scuffling; 4 pinning; 5 pinning and attempted biting). Male Long-Evans animals that were not included in the aggressor group were used as novel targets in the social interaction test.

#### Repeated Social Stressor

The bHR/bLR rats were randomly assigned to either the social defeat group (SD) or no defeat group (NIL). During the daily 15-minute social stress episode, Long-Evans females were removed from the resident cage and an intruder bHR or bLR rat (250 – 350g) was introduced. The animal was allowed to move freely throughout the cage until an aggressive interaction scoring at least 4 occurred. Following this, intruders were placed into a protective wire mesh cage (10 x 10 x 15cm) within the Long-Evans residents' home cage for the remainder of the trial. After the 15-minute social stress period, intruders were returned to their home cage with their cage-mates. All bHR/bLR intruders from one cage underwent social defeat at the same time. The female Long-Evans were returned to the cage of the resident aggressor at the end of each social defeat bout. Each bLR or bHR intruder was exposed to a different Long-Evans aggressor on each day of social defeat. Long-Evans aggressors were used multiple times in one social defeat session, with a minimum rest period of 40 minutes between each bout. Experimental bHR and bLR animals that did not experience the social stressor were placed in a clean, empty novel cage within the same testing room and allowed to move freely for the 15-minute period. Experimental bHR and bLR animals and Long-Evans resident aggressors were housed in separate rooms throughout the entirety of testing. Any bLR or bHR experimental subjects that sustained physical wounds during social defeat stress were excluded from the remainder of the study.

#### Testosterone, Oxytocin and Interleukin-6 ELISAs

Plasma testosterone and oxytocin levels were measured using enzyme immunoassay (EIA) kits from Arbor Assays (catalogue numbers K032 and K048, respectively https://www.arborassays.com/), and plasma IL-6 was measured using an EIA kit from BD Biosciences (catalogue number 550319 https://www.bdbiosciences.com/en-us).

Plasma for testosterone analysis underwent extraction prior to the EIA protocol; diethyl ether was added to plasma samples at a 5:1 ether:sample ratio, vortexed for two minutes and then allowed to rest for five minutes. Samples were then frozen in a dry ice bath and the top layer of diethyl ether pipetted off. This was then repeated, the two lots of ether from each sample combined, and then spun down in a speed-vacuum for 2 hours and stored at -20$^{o}$C overnight. Samples were re-dissolved at room temperature in 125μl of the assay buffer provided with the testosterone EIA kit and immediately processed.

Plasma for oxytocin analysis underwent extraction immediately prior to the EIA protocol; 100μl of plasma was mixed with 150μl of provided extraction solution, vortexed and shaken at room temperature for 90 minutes, then centrifuged at 1660 x g at 4$^{o}$C for 20 minutes. Supernatant was then drawn off and spun down in a speed-vacuum for 2 hours until dry. Samples were then reconstituted with 250μl of assay buffer and used in the EIA protocol. Plasma for IL-6 analysis did not undergo extraction or dissociation before use.

Following these preparatory steps, samples and the freshly prepared testosterone and oxytocin dilution standards were pipetted into well plates in duplicate, and in the case of IL-6 in triplicate. All provided protocols were then performed for the EIA kits, with a final step of 100μl of TMB substrate added to each well and incubated for 30 minutes at room temperature without shaking; 50μl of Stop Solution was then added to each well and optical density at 450nm of each well determined in a plate reader. Plasma concentrations of all substances were calculated using the Arbor Assays online calculator (https://www.myassays.com/).

#### Behavioral and Hormonal Data Analysis

##### Data and Code Availability

The full behavioral and hormonal dataset is available on Figshare (DOI: 10.6084/m9.figshare.24085524). All analyses were performed in Rstudio (v.1.0.153, R v. 3.4.1) (code: https://github.com/hagenaue/bHRbLR_Enrichment_Stress_BehaviorAndHormoneData).

##### Overview of Outcome Variables

We hypothesized that adolescent enrichment would influence the effects of social defeat on social and anxiety-related behaviors in a manner that depended on bred line. To test this hypothesis, our outcome variables included social behaviors on a social interaction test (social avoidance measured by Ethovision, social approach measured by Ethovision and video analysis, time on top measured by Ethovision and video analysis), exploration/anxiety-related behavior on the elevated plus maze (% time in the open arms), and exploration/anxiety-related behavior during habituation to the open field used for social interaction testing (distance traveled, % time in the center).

To properly interpret the effect of social defeat on these outcomes, we also examined the interactive effects of adolescent enrichment and bred line on behavior during social defeat (aggressive behavior, submissive behavior, social defeat score, time caged), as we hypothesized that each of these variables was likely to change the experience of social defeat itself.

We also hypothesized that adolescent enrichment would influence the effects of social defeat on the plasma levels of hormones important for mediating affective and social behaviors (corticosterone, testosterone, oxytocin, IL-6) in a manner that depended on bred line, although the results from the oxytocin and IL-6 assays ended up being uninterpretable due to many measurements overlapping the limits of sensitivity for the assays. Altogether, our study considered 16 behavioral and hormonal outcome variables of interest, followed by an exploratory RNA-Seq analysis performed on nucleus accumbens (NACC) and hippocampal (HC) tissue collected at the time of sacrifice.

##### Experimental Design

Sample sizes varied across behavioral tasks and biological measurements (**Fig S1**), with full sample sizes ranging from *N*=25-142, and subgroup sample sizes ranging from *n*=5-14/subgroup. The sample sizes were determined by animal availability, budget, and logistical capacity. By rough estimation (http://fpr-calc.ucl.ac.uk), within the full sample size (n=142), there was sufficient power (80%) to reliably detect at a traditional alpha ($p<0.05$) medium effect sizes for the main effects for each of the variables of interest (Line: Cohen's d>0.47, Social Defeat: d>0.47, Enrichment: d>0.51) and their second order interactions (d>0.67) and large effect sizes for their third order interactions (d>0.98).

**Full Sample Size:**

| | bHR NIL | bHR SE | bHR EE | bHR NIL + SD | bHR SE + SD | bHR EE + SD | bLR NIL | bLR SE | bLR EE | bLR NIL + SD | bLR SE + SD | bLR EE + SD |
|---|---|---|---|---|---|---|---|---|---|---|---|---|
| F49 | 6 | 7 | 6 | 4 | 4 | 6 | 2 | 6 | 6 | 4 | 5 | 6 |
| F53 | 0 | 3 | 3 | 0 | 3 | 3 | 3 | 3 | 3 | 0 | 3 | 3 |
| F56 | 6 | 3 | 3 | 8 | 3 | 3 | 5 | 2 | 4 | 8 | 3 | 5 |

***Colored to illustrate bHR and bLR sample size variation across subgroups***

**Variables Measured:**

***Elevated Plus Maze: Time in the Open Arms, Open Field Habituation: Distance Traveled***

**Social Interaction Test:**

| | bHR NIL | bHR SE | bHR EE | bHR NIL + SD | bHR SE + SD | bHR EE + SD | bLR NIL | bLR SE | bLR EE | bLR NIL + SD | bLR SE + SD | bLR EE + SD |
|---|---|---|---|---|---|---|---|---|---|---|---|---|
| F49 | 6 | 7 | 6 | 4 | 4 | 6 | 2 | 6 | 6 | 4 | 5 | 6 |
| F53 | 0 | 3 | 3 | 0 | 3 | 3 | 3 | 3 | 3 | 0 | 3 | 3 |
| F56 | 6 | 3 | 3 | 8 | 3 | 3 | 5 | 2 | 4 | 8 | 3 | 5 |

**Variables Measured:**

***Social Interaction, Social Avoidance, Time on Top of Stimulus Animal (Scored by Ethovision)***

***Social Interaction, Social Avoidance, Time on Top of Stimulus Animal (Scored by Ethovision & Hand)***

**Social Defeat Sessions: Days 1-4**

| | bHR NIL | bHR SE | bHR EE | bHR NIL + SD | bHR SE + SD | bHR EE + SD | bLR NIL | bLR SE | bLR EE | bLR NIL + SD | bLR SE + SD | bLR EE + SD |
|---|---|---|---|---|---|---|---|---|---|---|---|---|
| F49 | 6 | 7 | 6 | 4 | 4 | 6 | 2 | 6 | 6 | 4 | 4 | 6 |
| F53 | 0 | 3 | 3 | 0 | 3 | 3 | 3 | 3 | 3 | 0 | 3 | 3 |
| F56 | 6 | 3 | 3 | 8 | 3 | 3 | 5 | 2 | 4 | 8 | 3 | 5 |

**Variables Measured:**

***Social Defeat Score, Time Caged with Aggressor, Video Analysis: Submissive, Aggressive and Other Behaviors***

**Hormone Measurements at Time of Sacrifice**

| | bHR NIL | bHR SE | bHR EE | bHR NIL + SD | bHR SE + SD | bHR EE + SD | bLR NIL | bLR SE | bLR EE | bLR NIL + SD | bLR SE + SD | bLR EE + SD |
|---|---|---|---|---|---|---|---|---|---|---|---|---|
| F49 | 6 | 7 | 6 | 4 | 4 | 6 | 2 | 6 | 6 | 4 | 5 | 6 |
| F53 | 0 | 3 | 3 | 0 | 3 | 3 | 3 | 3 | 3 | 0 | 3 | 3 |
| F56 | 6 | 3 | 3 | 8 | 3 | 3 | 5 | 2 | 4 | 8 | 3 | 5 |

**Variables Measured:**

**Corticosterone, Testosterone, IL6**

**Corticosterone, Testosterone, IL6, Oxytocin**

**Brain Tissue: Hippocampal RNA-Seq**

| | bHR NIL | bHR SE | bHR EE | bHR NIL + SD | bHR SE + SD | bHR EE + SD | bLR NIL | bLR SE | bLR EE | bLR NIL + SD | bLR SE + SD | bLR EE + SD |
|---|---|---|---|---|---|---|---|---|---|---|---|---|
| F49 | 6 | 7 | 6 | 4 | 4 | 6 | 2 | 6 | 6 | 4 | 5 | 6 |
| F53 | 0 | 3 | 3 | 0 | 3 | 3 | 3 | 3 | 3 | 0 | 3 | 2 (3) |
| F56 | 6 | 3 | 3 | 8 | 3 | 3 | 5 | 2 | (2) 4 | 8 | 3 | 5 |

**Brain Tissue: NAcc RNA-Seq**

| | bHR NIL | bHR SE | bHR EE | bHR NIL + SD | bHR SE + SD | bHR EE + SD | bLR NIL | bLR SE | bLR EE | bLR NIL + SD | bLR SE + SD | bLR EE + SD |
|---|---|---|---|---|---|---|---|---|---|---|---|---|
| F49 | 6 | 7 | 6 | 4 | 4 | 6 | 2 | 6 | 6 | 4 | 5 | 5 (6) |
| F53 | 0 | 3 | 3 | 0 | 3 | 3 | 3 | 3 | 3 | 0 | 3 | 3 |
| F56 | 6 | 3 | 3 | 8 | 3 | 3 | 5 | 2 | 4 | 8 | 3 | 3 (5) |

##### *Fig S 1. The subgroup sample sizes associated with each dependent variable.*

*Within the top table (Full sample size), color is used to illustrate subgroups with more subjects (darker) versus fewer subjects (lighter), with the hue of the color illustrating bHR (green) versus bLR (red) groups. In other panels, color is used to illustrate which subgroups were used for each set of related variables. With the exception of the F56 bHR NIL and F56 bHR + SD animals, all animals from a particular generation within a particular subgroup are from the same litter (i.e., each cell in the table typically represents animals from the same litter). Numbers in parentheses indicate the original sample for the RNA-Seq data vs. the final sample following quality control.*

Due to logistics and the need to keep the study design manageable, some variables were restricted in terms of generation or experimental subgroups represented. Hand-scored social interaction behaviors were only available from generations F53 and F56. That sample size (total n=80) had sufficient power (80%) to reliably detect at a traditional alpha (p<0.05) medium effect sizes for the main effects for each of the variables of interest (Line: d>0.64, Social Defeat: d>0.64, Enrichment: Cohen's d>0.66), large effect sizes for their second order interactions (d>0.91), and very large effect sizes for their third order interactions (d>1.32).

Hormone measurements were only available from generations F53 and F56 and were restricted to standard housing (NIL) and enhanced (EE) subgroups. That sample size (total n=57) had sufficient power (80%) to reliably detect at a traditional alpha (p<0.05) large effect sizes for the main effects for each of the variables of interest (Cohen's d>0.76 for Line, Enrichment, and Social Defeat) and for their second order interactions (d>1.08), and very large effect sizes for their third order interactions (d>1.57).

RNA-Seq data was only collected from bLR rats in the standard housing (NIL) and enhanced enrichment (EE) subgroups. For the NACC, the sample size (total n=46) had sufficient power (80%) to reliably detect at a traditional alpha (p<0.05) large effect sizes for the main effects for each of the variables of interest (Cohen's d>0.84 for Enrichment and Social Defeat) and very large effect sizes for their second order interaction (d>1.23). The HC was more weakly powered: the sample size (total n=25) had sufficient power (80%) to reliably detect very large effect sizes for the main effects for each of the variables of interest (Enrichment: Cohen's d>1.17, Social Defeat: d>1.19) and their second order interaction (d>1.8).

Social defeat related variables were measured for animals that were socially-defeated (SD) across all four days of defeat (*i.e.,* a within-subjects time series), a few animals (6-17 out of 71) missed measurements on any particular day due to technical issues (**Fig S2**). That sample size (n=70 subjects) had sufficient power (80%) to reliably detect at a traditional alpha (p<0.05) large effect sizes for the main effects for each of the between-subject variables of interest (Line: Cohen's d>0.68, Enrichment: d>0.72) and second order interaction (d>0.98).

**Social Defeat Day 1**

| | bHR NIL | bHR SE | bHR EE | bHR NIL + SD | bHR SE + SD | bHR EE + SD | bLR NIL | bLR SE | bLR EE | bLR NIL + SD | bLR SE + SD | bLR EE + SD |
|---|---|---|---|---|---|---|---|---|---|---|---|---|
| F49 | 6 | 7 | 6 | 4 | 4 | 6 | 2 | 6 | 6 | 4 | 4 | 6 |
| F53 | 0 | 3 | 3 | 0 | 3 | 3 | 3 | 3 | 3 | 0 | 3 | 3 |
| F56 | 6 | 3 | 3 | 8 | 3 | 3 | 5 | 2 | 4 | 8 | 3 | 0 |

**Social Defeat Day 2**

| | bHR NIL | bHR SE | bHR EE | bHR NIL + SD | bHR SE + SD | bHR EE + SD | bLR NIL | bLR SE | bLR EE | bLR NIL + SD | bLR SE + SD | bLR EE + SD |
|---|---|---|---|---|---|---|---|---|---|---|---|---|
| F49 | 6 | 7 | 6 | 4 | 4 | 6 | 2 | 6 | 6 | 4 | 0 | 6 |
| F53 | 0 | 3 | 3 | 0 | 3 | 3 | 3 | 3 | 3 | 0 | 3 | 3 |
| F56 | 6 | 3 | 3 | 7 | 3 | 3 | 5 | 2 | 4 | 7 | 3 | 5 |

**Social Defeat Day 3**

| | bHR NIL | bHR SE | bHR EE | bHR NIL + SD | bHR SE + SD | bHR EE + SD | bLR NIL | bLR SE | bLR EE | bLR NIL + SD | bLR SE + SD | bLR EE + SD |
|---|---|---|---|---|---|---|---|---|---|---|---|---|
| F49 | 6 | 7 | 6 | 4 | 4 | 0 | 2 | 6 | 6 | 4 | 4 | 0 |
| F53 | 0 | 3 | 3 | 0 | 3 | 3 | 3 | 3 | 3 | 0 | 3 | 3 |
| F56 | 6 | 3 | 3 | 5 | 2 | 3 | 5 | 2 | 4 | 8 | 3 | 5 |

**Social Defeat Day 4**

| | bHR NIL | bHR SE | bHR EE | bHR NIL + SD | bHR SE + SD | bHR EE + SD | bLR NIL | bLR SE | bLR EE | bLR NIL + SD | bLR SE + SD | bLR EE + SD |
|---|---|---|---|---|---|---|---|---|---|---|---|---|
| F49 | 6 | 7 | 6 | 0 | 4 | 6 | 2 | 6 | 6 | 0 | 4 | 6 |
| F53 | 0 | 3 | 3 | 0 | 3 | 3 | 3 | 3 | 3 | 0 | 3 | 3 |
| F56 | 6 | 3 | 3 | 8 | 3 | 3 | 5 | 2 | 4 | 7 | 3 | 5 |

**Variables Measured:**

***Video Analysis: Submissive, Aggressive and Other Behaviors***

***Note: Social Defeat Score and Time Caged with Aggressor are still available for days when video footage is missing***

***Fig S 2. The subgroup sample sizes with video footage for each day of social defeat.***

*Color is used to illustrate subgroups with more subjects (red) versus fewer subjects (white).*

#### Data Distribution

Many of the variables yielded non-normally distributed data (**Fig S3**) and were likely to be drawn from a non-normally distributed population (bounded data-types). Behavioral variables were often defined as percentages of the total time spent on a task, and thus were bounded between 0-100% with skewed or otherwise non-normal distributions and common outliers. Hormone measurements were bounded at 0 (or, more realistically, at the limit of detection for the assay), and highly skewed in the positive direction. These non-normal distributions indicated that inferential statistics were best performed following data transformation or using non-parametric statistical methods.

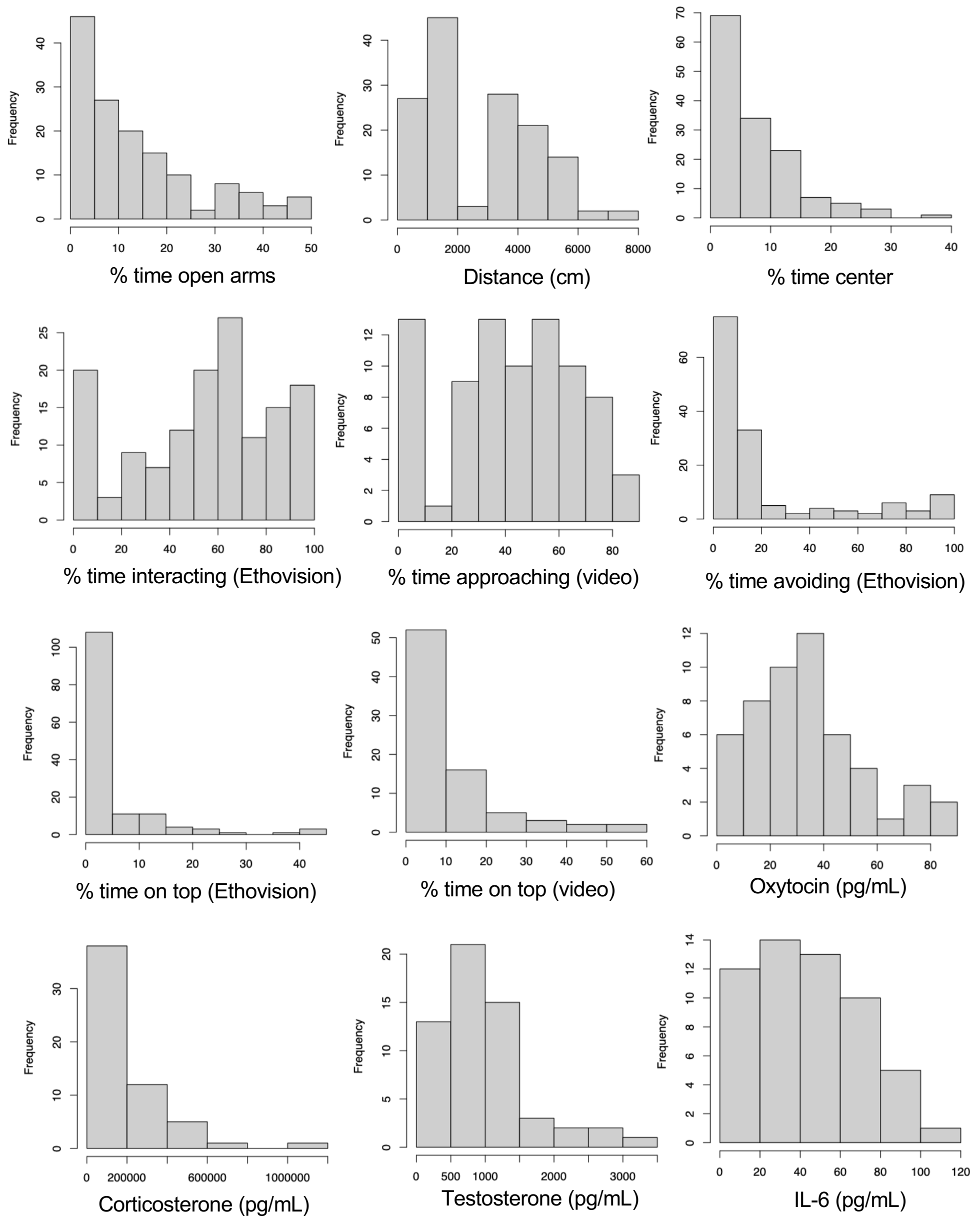

***Fig S 3. Histograms showing that most of the dependent variables had a non-normal distribution.***

These non-normal distributions indicate that inferential statistics are best performed following transformation or using non-parametric statistical methods. Behavioral variables were measured using the elevated plus maze (% time open arms), novel open field (distance, % time center), and social interaction task (% time interacting, % time approaching, % time avoiding, % time on top).

#### Statistical Tests

The statistical tests performed to assess the effect of bred line, adolescent enrichment, and social defeat on each of our behavioral and hormonal outcome variables were chosen by a professional analyst following data collection and a review of the experimental design and data distributions.

Due to the non-normally distributed data and small subgroup sample sizes, we primarily used non-parametric statistical tests to evaluate our findings. We also included generation (F49 vs. F53 vs. F56) as a categorical co-variate to control for batch effects, as the animals from each generation went through enrichment, social defeat, behavioral testing, sacrifice, blood collection, and brain extraction in tandem.

To examine the influence of categorical predictors bred line (bHR vs. bLR), adolescent enrichment (NIL vs. SE vs. EE), social defeat (NIL vs. SD) on continuous variables, we used permutation-based linear regression or ANOVA using the functions *lmperm()* and *aovperm()* available in the R package *permuco* [3] with 15,000 permutations (alpha=0.05). Permutation-based methods evaluate effects in data without making strong assumptions regarding the shape of the sampling distribution, although the tests still assume exchangeability – *i.e.,* that the distribution for our subgroups would be identical if the null hypothesis were true, with similar variance and skew. We used the method of permutation proposed by Freedman and Lane [4], which permutates residuals derived from a reduced model (containing only the nuisance variables). These residuals are then added to the coefficients derived from a full model run on the raw data, and the full model is then re-run to produce an empirical distribution for the T-statistic. This method has been shown to produce better control of Type 1 error in multi-factorial linear regression and ANOVA-style analyses performed on data with small sample sizes, non-normally distributed residuals, and outliers, while simultaneously improving power by 3-9% as compared to traditional parametric tests [5-7].

Our design was occasionally unbalanced, which can introduce moderate multicollinearity amongst our predictor variables and affect the interpretation of model coefficients. Therefore, for all linear regression and ANOVA-style tests, we used Type III (“unique”, “orthogonal”, or “marginal”) Sum of Squares, which produces results for each variable while controlling for the effects of all other variables (as if the variable was entered last into the model). For the linear regression-style tests, we used contrasts defined by treatment (*i.e.,* the difference between each level of a factor and the reference level), with the reference levels for each variable defined as bHR (bred line), NIL (no adolescent enrichment), NIL (no social defeat), and F53 (generation). For ANOVA-style methods, we used contrasts defined by sum, so that the intercept was centered on the weighted mean between the conditions. For this output, the effect of any particular variable can be considered as a representation of the general effect across conditions (*e.g.,* the effect of social defeat on a hypothetical "average" animal in our experiment).

Permutation-based methods are less well-equipped to handle the violations of sphericity introduced by time-series analyses. Therefore, for the analysis of the four days of social defeat-related behavior, we used a multilevel model that included a linear effect of time (the day of social defeat) and ratID as a random effect using the function *lme()* from the R package *nlme* [8] an autocorrelation structure of order 1 (AR1), and a model fit by maximizing log-likelihood ("ML"). To satisfy the assumption of normality for this test, we applied a log2 transformation to the percent time spent in aggressive and submissive behavior. Within this model, we examined the interaction of the effects of the day of social defeat with bred line and adolescent enrichment, using generation as a co-variate:

*y~Day*Line*Enrichment+Generation, random=~Day|RatID*

To make the interpretation of the results more comparable to later behavioral tests, the intercept for the model was defined as Day 4 of social defeat, so that the coefficients for bred line and adolescent enrichment would reflect the cumulative effects of all four days of defeat. The effects of the factors in this model were then summarized using the *Anova()* function from the R package *car* [9], using type III Sum of Squares and contrasts defined by sum, and significance was determined using a likelihood ratio test.

Due to the large number of behavioral and hormonal dependent variables (16), a Bonferonni-corrected alpha (adj.p<0.05) defined significance (denoted *). Results at a traditional alpha (p<0.05) are called nominal (denoted #) in the text and considered tentative.

#### RNA-Seq Data Analysis

##### Data and Code Availability

The full nucleus accumbens (NACC) and hippocampus (HC) RNA-Seq datasets can be found on GEO/SRA at (Accession# GSE237890, https://www.ncbi.nlm.nih.gov/geo/query/acc.cgi?acc=GSE237890). The initial RNA-Seq data preprocessing was performed using a standardized pipeline within the MBNI Analysis Hub ("Ahub": https://ahub.mbni.org). All later downstream analyses were performed in Rstudio (v.1.0.153, R v. 3.4.1), with full code available at https://github.com/hagenaue/bHRbLR_Enrichment_Stress_RNASeqData.

##### Alignment and Assembly

Within the Ahub pipeline, RNA-Seq data was aligned using the STAR algorithm (2.7.0f, Encode parameter: unique hit only) to genome assembly Rnor6. Reads were then summarized into counts per transcript using featureCounts (v1.6.4) and Ensembl 96 annotation.

#### Quality Control

Within the Ahub pipeline, principal components analysis was used to identify extreme outlier samples, which were presumed to reflect unknown sources of technical variation (NACC: 2 samples, HC: 6 samples). These samples did not represent any particular treatment group or processing batch and were removed before further analysis was performed.

Additional quality control and normalization was performed in Rstudio (v.1.0.153, R v.3.4.1) for the NACC and HC datasets independently. Within the NACC dataset, library size was initially found to vary from 1.5 million to 53 million (median=30 million, interquartile range: 22-33 million). The initial count matrix included reads aligned to 19,209 Ensembl-annotated transcripts (rows with reads>0). This matrix was filtered to remove transcripts with low-level expression (an average of less than 1 read across all subjects), leaving data from 17,765 transcripts. To correct for artifacts introduced by sample-level differences in RNA production, normalization factors were calculated using the trimmed mean of M-values (TMM) method [10] within the *calcNormFactors()* function (package: *edgeR, v.3.18.1*) and then the data was transformed to Log2 counts per million (cpm) [11]. One sample had a notably low distribution of log2 cpm and was removed, leaving a final sample size of 46 subjects (bLR NIL: n=10, bLR NIL+SD: n=12, bLR EE: n=13, bLR EE+SD: n=11).

Within the HC dataset, library size was initially found to vary from 1.8 million to 47 million (median=20 million, interquartile range: 10-35 million). The initial count matrix included reads aligned to 19,045 Ensembl-annotated transcripts (rows with reads>0). This matrix was filtered to remove transcripts with low-level expression (an average of less than 1 read across all subjects), leaving data from 17,629 transcripts. Following TMM normalization, the data was transformed to Log2 cpm. The final sample size was 25 subjects (bLR NIL: n=5, bLR NIL+SD: n=8, bLR EE: n=5, bLR EE+SD: n=7).

#### Model Selection

To determine which technical variables might be important co-variates to include in the differential expression model, we first evaluated the quality-controlled dataset for confounding collinearity between treatment and technical variables using either Fisher's exact test (categorical vs. categorical), linear regression (numeric vs. categorical), or one-way ANOVA (numeric vs. categorical). Technical variables were then further evaluated for potential impact on differential expression results by examining their relationship with the top principal components of variation in the data (PC1-PC3, as calculated using log2 cpm data that had been centered and scaled by gene).

Within the NACC dataset, there was a negative relationship between Enrichment and sample library size ($\beta$=-10,927,086, p=6.78e-05, **FigS4**) and an independent trend towards a negative relationship with RNA concentration (β=-14.135, p=0.0697). There was also a trend towards a relationship between Social Defeat and Date of Dissection (Fisher's Exact Test: p=0.07168). Generation, RNA Purity (260/280), Date of RNA Extraction, and Hemisphere Dissected were not unevenly distributed in relationship to treatment group (EE or SD, *p*>0.10).

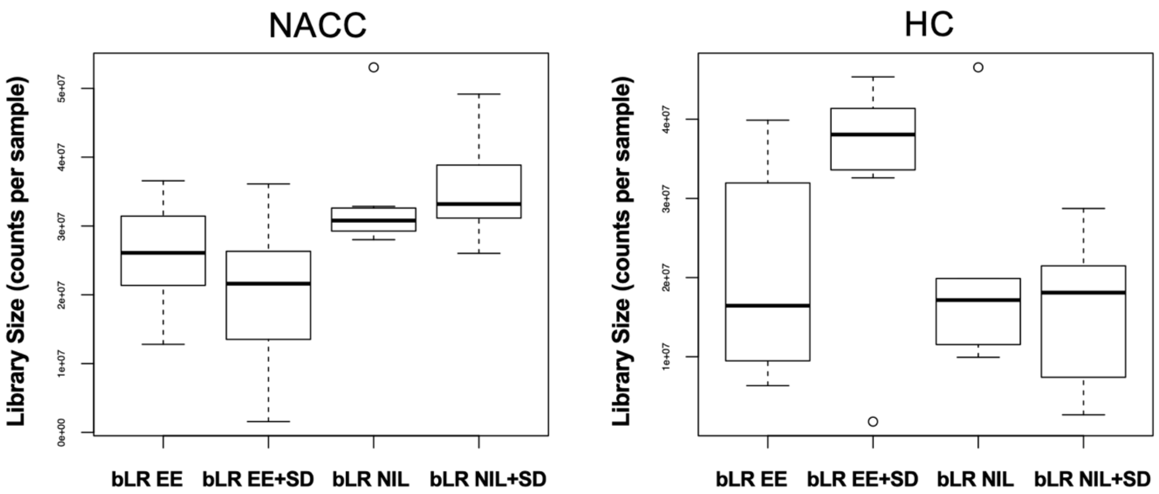

***Fig S 4. RNA-Seq quality control: Library size varies with treatment group.***

*In both the NACC and HC RNA-Seq datasets, library size was found to vary with treatment group, but in a manner that differed by brain region. In the NACC, library size was lower following EE (p=6.78e-05), whereas in the HC, library size showed a trend towards being greater following EE (p=0.0636). This variation could be due to biological or technical factors, but did not appear to be an important source of variation in the normalized data, as indicated by principal components analysis.*

Within the NACC, PC1 accounted for 16% of the variation in the dataset (**Fig S5**) and was related to RNA concentration (p=0.0195) and Date of Dissection (ANOVA: p=0.09582*trend, *post-hoc*: p=0.0418 for 17/1/2019 in particular), and showed a trend towards a relationship with Social Defeat (p=0.08033). PC2 accounted for 13% of the variation in the dataset and was related to Date of RNA extraction (p=0.003305), Generation (p=0.00212), and Date of dissection (ANOVA: p=0.00204, particularly driven by 21/2/2019: p=0.000497), as well as a trend towards a relationship with Enrichment (p=0.087886). Library size did not seem to be strongly related to any of the top principal components of variation in the dataset, probably due to normalization**.** Based on these results, the co-variates that were deemed most likely to be important were RNA concentration, RNA extraction Batch, Dissection Date Batches and Generation. Since the Dissection Date Batches and Generation closely overlapped, we chose to include Dissection Date as the co-variate in our models, because it was more collinear with Social Defeat and preliminary analyses indicated that it was also more strongly related to gene expression.

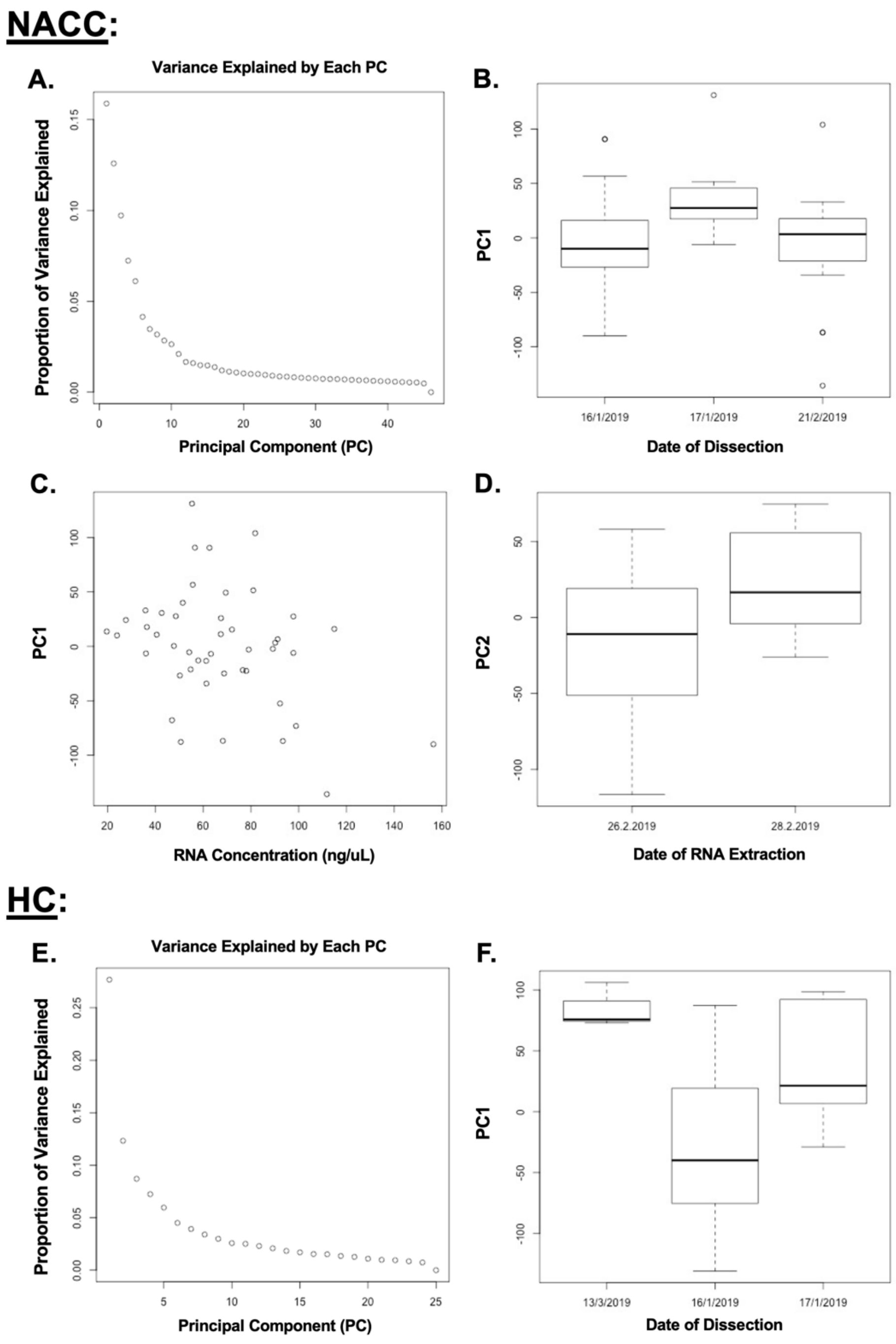

***Fig S 5. RNA-Seq quality control: Identifying important technical covariates for the differential expression model using principal components analysis.***

***A)** A scree plot illustrating the proportion of variance explained by each principal component of variation in the NACC dataset. **B)** Boxplot showing the relationship between PC1 and Date of Dissection (p=0.0418), **C**) Scatterplot showing the correlation between PC1 and RNA concentration (p=0.0195) **D**) Boxplot showing the correlation between PC2 and Date of RNA extraction (p=0.003305). **E)** A scree plot illustrating the proportion of variance explained by each principal component of variation in the HC dataset. **F)** A boxplot illustrating the relationship between PC1 and Date of Dissection (p=0.00687).*

Within the HC dataset, there was a relationship between Enrichment and Generation (Fisher's Exact Test: p=0.01491, **Fig S1**) and a trend towards a positive relationship between Enrichment and sample library size (β=10,659,538, p=0.06359, **Fig S4**) and between Social Defeat and Date of Dissection (Fisher's Exact Test: p=0.0645). RNA Concentration, RNA Purity (260/280), Date of RNA Extraction, and Hemisphere Dissected were not unevenly distributed in relationship to treatment group (EE or SD, $p$>0.10).

Within the HC, PC1 accounted for 28% of the variation in the dataset and was related to Date of Dissection (ANOVA: p= 0.006866, **Fig S5**). PC2 accounted for 13% of the variation in the dataset, and was related to library size (R=0.69). Based on these results, Date of Dissection and Library Size were prioritized as potentially important co-variates, but our sample size was relatively small and preliminary analyses indicated that Library Size was not strongly related to gene expression when included in a model with SD, EE, and Date of Dissection (no transcripts with FDR<0.10), so we chose a final model with only the co-variate of Date of Dissection.

#### Differential Expression Analysis

Differential expression was calculated using the limma/voom method ([12] package: *limma* v.3.32.5, [13]) with observed precision weights in a weighted least squares linear regression using the following models:

##### NACC:

*Model 1 ("M1: Main Effects Model"):*

$$y \sim \beta_0 + \beta_1 SocialDefeat + \beta_2 Enrichment + \beta_3 RNAconc + \beta_4 RNAextractBatch + \beta_{5-6} DissectionBatches + \varepsilon$$

*Model 2 ("M2: Interactive Effects Model"):*

$$y \sim \beta_0 + \beta_1 SocialDefeat + \beta_2 Enrichment + \beta_3 (SocialDefeat * Enrichment) + \beta_4 RNAconc + \beta_5 RNAextractBatch + \beta_{6-7} DissectionBatches + \varepsilon$$

### HC:

*Model 1 ("M1: Main Effects Model"):*

$$y \sim \beta_0 + \beta_1 SocialDefeat + \beta_2 Enrichment + \beta_{3-4} DissectionBatches + \varepsilon$$

*Model 2 ("M2: Interactive Effects Model"):*

$$y \sim \beta_0 + \beta_1 SocialDefeat + \beta_2 Enrichment + \beta_3 (SocialDefeat * Enrichment) + \beta_{4-5} DissectionBatches + \varepsilon$$

Contrasts were defined by treatment, with the intercept for Enrichment and Social Defeat both set as “NIL”. Standard error was then moderated towards a global value provided by an empirical Bayes distribution (function *eBayes())* and multiple comparison correction was performed using the Benjamini-Hochberg method (false discovery rate (FDR) or q-value, [14]).

#### Sensitivity of Results to Model Specification

Following these results, the results of reduced models were explored to determine the sensitivity of our findings to model specification. Within the NACC dataset, all models produced a very similar list of top genes for both adolescent enrichment and social defeat, with the top enrichment genes dominated by protocadherins and the top social defeat genes dominated by RT1 related genes. In general, the coefficients for social defeat or enrichment produced by the M1 model with or without co-variates were highly correlated (EE: $R^2$=0.84, SD: $R^2$=0.62). The coefficients for social defeat or enrichment produced by the M2 model with or without co-variates typically had greater effect sizes and were slightly less correlated (EE: $R^2$=0.53, SD: $R^2$=0.62). In the regression models that included an interactive effects term, there was a slight correlation between the main effects of EE and SD ($R^2$=0.14), that could be due to their shared control group (bLR NIL NIL), that disappeared when the model only included the main effects of EE and SD ($R^2$=0). Therefore, we chose to emphasize findings that were evident in the results from both M1 and M2. Within the small HC dataset, models with fewer terms were better powered to detect main effects of EE, whereas none of the models showed significant effects of SD or SD*EE (FDR>0.1).

#### Functional Ontology

We evaluated whether our differential expression results were enriched for genes representing particular functional, anatomical, and cell-type categories using fGSEA [15] (v.1.2.1, nperm=10000, minSize = 10, maxSize = 1000) and a custom gene set file (Brain.GMT v.2 for rats, # of gene sets: 15545). To run the analysis, we used differential expression results that were annotated by official gene symbol and pre-ranked by either log2FC or T-stat, with any results that mapped to the same gene symbol averaged. The log2FC version of the analysis seemed to be biased in favor of low-expressed genes, which tended to show more extreme effect sizes. Therefore, the T-stat version of the analysis was chosen for the final result.

The custom gene set file included gene sets from three commonly-used collections on the Molecular Signatures Database (MSigDB v7.3, http://software.broadinstitute.org/gsea/msigdb/index.jsp, downloaded 2021-03-25): the full traditional Gene Ontology annotation database (“C5: GO Gene Sets”, file: “c5.all.v7.3.symbols.gmt.txt”, # of gene sets: 14,996), as well as two other databases that had been subsequently filtered for relevance to central nervous system tissue: the curated gene set database (“C2: Curated Gene Sets”, file: “c2.all.v7.3.symbols.gmt.txt”, # of filtered gene sets: 158) and the cell type database (“C8: Cell Type Signature Gene Sets”, file: “c8.all.v7.3.symbols.gmt.txt”, # of filtered gene sets: 211).

The custom gene set file also included gene sets enriched for expression within specific brain cell types from additional databases (BrainInABlender database [16]: *https://github.com/hagenaue/BrainInABlender*, v.0.0.0.9000, # of gene sets: 39); DropViz scRNA-Seq database: [17], *http://dropviz.org,* HC and striatal gene sets (# of gene sets: 25), and gene sets curated ([18], # of gene sets: 69) to target HC regional gene expression signatures [19] and HC co-expression networks [20, 21].

The custom gene set file also included gene sets designed to provide insight into the role of the NACC and HC in processing affective behavior. Some of these gene sets were designed to allow us to quickly and uniformly assess the overlap of our findings with the lists of differentially expressed genes identified in related brain transcriptional profiling publications, including those examining the effects of stress in the NACC and HC [22-25] (# of gene sets: 14), the effects of selective breeding targeting internalizing behavior in the HC (curated in [18], # of gene sets: 19), and the effects of human neuropsychiatric disorders in the cortex ([26]: # of gene sets: 14). Then, to gain a more comprehensive comparison, we extracted the differential expression results from all of the studies in the NACC or HC related to stress, enrichment, affective behavior, and mood disorder from two online databases of differential expression results: Gemma (https://gemma.msl.ubc.ca/home.html, [27]) and GeneWeaver (https://www.geneweaver.org, [28]) (GeneWeaver: n=38 gene sets, NACC=5, HC=33; Gemma: n=329 gene sets, NACC: 28, HC: 301).

Since our custom gene set database included many gene sets from each of these categories, we were more likely to see enrichment within gene sets in these categories simply due to random chance. Therefore, we assessed whether there was a *disproportionate* enrichment within each category in our results. The gene sets in the custom gene set file related to social stress, fear conditioning, other stress, corticosterone, enrichment, social & aggressive behavior, locomotor activity, mood, and fear/anxiety were identified by an experimenter blinded to the fGSEA results. To determine whether there was disproportionate enrichment within these categories (**Fig 5**), these results were then extracted and the percent of gene sets significantly enriched with differential expression (FDR<0.05 for M1 or M2 results) in each category was quantified and compared to the percent of gene sets enriched for differential expression (FDR<0.05 for M1 or M2 results) in the full fGSEA output (minus the category of interest) using Fisher’s Exact Test. A Bonferonni-corrected alpha (36 comparisons, adj.p<0.05) was used to define significance (denoted *). Results at a traditional alpha (p<0.05) are called nominal in the text (denoted #) and considered tentative.

### Supplementary Results

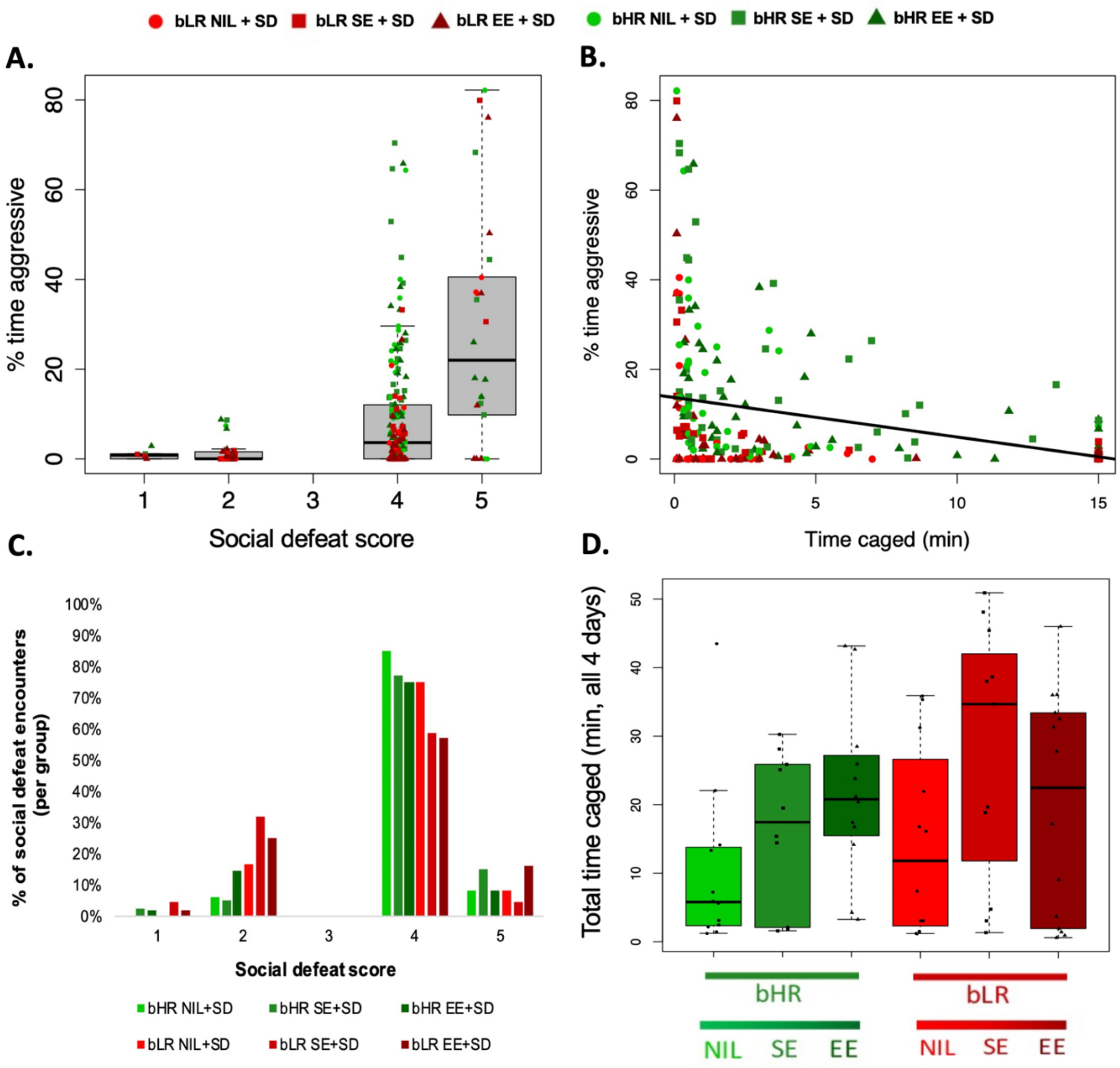

***Fig S 6. The resident Long-Evans aggressor animals appeared to respond to aggressive behavior during social defeat sessions with more severe defeat (social defeat scores>4), leading to a nominally earlier intervention by the experimenter to stop direct interaction (time caged lower than the full 15 minutes).***

*The more aggressive bHR animals typically experienced more severe defeat and less time caged with the aggressor. Bred line is illustrated with box fill color (bHR=green, bLR=red), adolescent enrichment by datapoint shape (circle=standard housing (NIL), square=simple enrichment (SE), triangle=enhanced enrichment (EE)).* ***A.*** *The percent time that the intruder bHR or bLR rats spent engaging in aggressive behavior during the four social defeat sessions was associated with more severe social defeat scores (1=lowest severity, 5=highest severity).* ***B.*** *The percent time that the intruder bHR or bLR rats spent engaging in aggressive behavior during the four social defeat sessions was associated with less time spent directly caged with* *the Long-Evans aggressor (minutes, maximum=15). **C.** Intruder bHR rats experienced a greater percent of encounters with more severe social defeat (greater social defeat score) (full time series not shown). **D.** Intruder bLR rats experienced a nominally greater duration of time (min) spent caged with the Long-Evans aggressor over the four defeat sessions (four 15 min sessions, or 60 min total) (full time series not shown). Boxplots illustrate the median and interquartile range for each treatment group (+/- whiskers illustrating the range and/or 1.5x interquartile range.*

#### Exposure to adolescent enrichment altered bLR and bHR social behavior

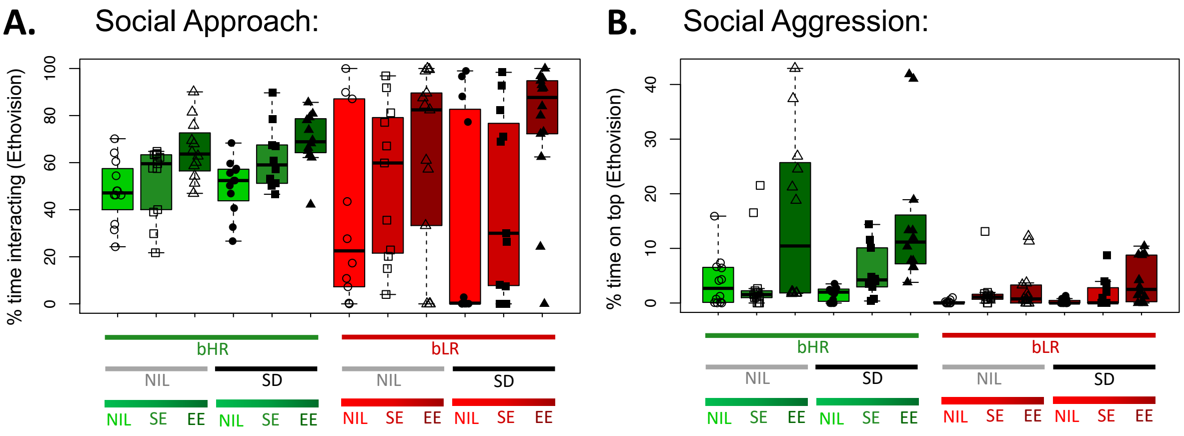

***Fig S 7. Adolescent social and environmental enrichment increased approach behavior in bLRs and increased aggression in bHRs: automated Ethovision analysis.***

***A-B.** Boxplots illustrate the median and interquartile range for each treatment group (+/- whiskers illustrating the range and/or 1.5x interquartile range). Bred line is illustrated with box fill color (bHR=green, bLR=red), adolescent enrichment by datapoint shape (circle=standard housing (NIL), square=simple enrichment (SE), triangle=enhanced enrichment (EE)), and social defeat is indicated by datapoint color (open=no defeat (NIL), black filled=defeated (SD)). **A.** Adolescent enrichment increased the percent time spent in the social interaction zone, especially for bLRs, as measured by automated Ethovision analysis. **B.** Adolescent enrichment increased the percent time on top of the stimulus animal's cage, especially for more aggressive bHRs, as measured by automated Ethovision analysis.*

#### Open-field center exploration revealed expected bHR/bLR differences, but showed an unexpected pattern following enrichment and defeat

Previous work has demonstrated that bHR and bLR animals behave differently within the open-field arena, with bHR animals spending more time in the center of the open-field [29]. Within our study, we similarly observed that the amount of time spent exploring the center of the open field was strongly influenced by bHR/bLR phenotype (**Fig S8**, main effect of Line: F(1,128)=49.40, p=6.667e-05*), but we also observed an unexpected nominal decrease in response to enrichment (main effect of enrichment: F(2,128)=5.72, p=3.667e-03#), that was larger in bHRs (interactive effects of Line*Enrichment: F(2, 128)=10.31, p=6.667e-05*), as well as an unexpected increase in response to social defeat (main effect of Social Defeat: F(1,128)=22.16, p=6.667e-05*). Traditionally, the amount of time exploring the center of the open field correlates with the results of other tasks contrasting the motivation to explore with anxiety (such as the elevated plus maze). However, in our experiment the open-field task was used to habituate animals to the testing arena prior to social interaction testing; therefore animals were placed into the center of the arena to prevent side preferences from developing. This often resulted in animals freezing within the center of the open-field, thus preventing traditional interpretation of open-field data. This minimal impact of adolescent enrichment on open-field behaviour aligns with previous observations in outbred Sprague-Dawley animals [30, 31], and provides further evidence that rodents exposed to enriched environments may adapt their behaviour during testing in unexpected ways due to an altered reaction to novel environments [32]. Possibly, a greater impact of enrichment in bHR rats was observed in the EPM than the open-field due to animals' experience of the multi-level enriching cage having similar ledges and height as the EPM maze. Interactions between genetic background and life experience resulted in variable reactions to different behavioural testing apparatus, suggesting that running a battery of tests enables greater insight into animal behaviours than just one behavioural assessment would provide.

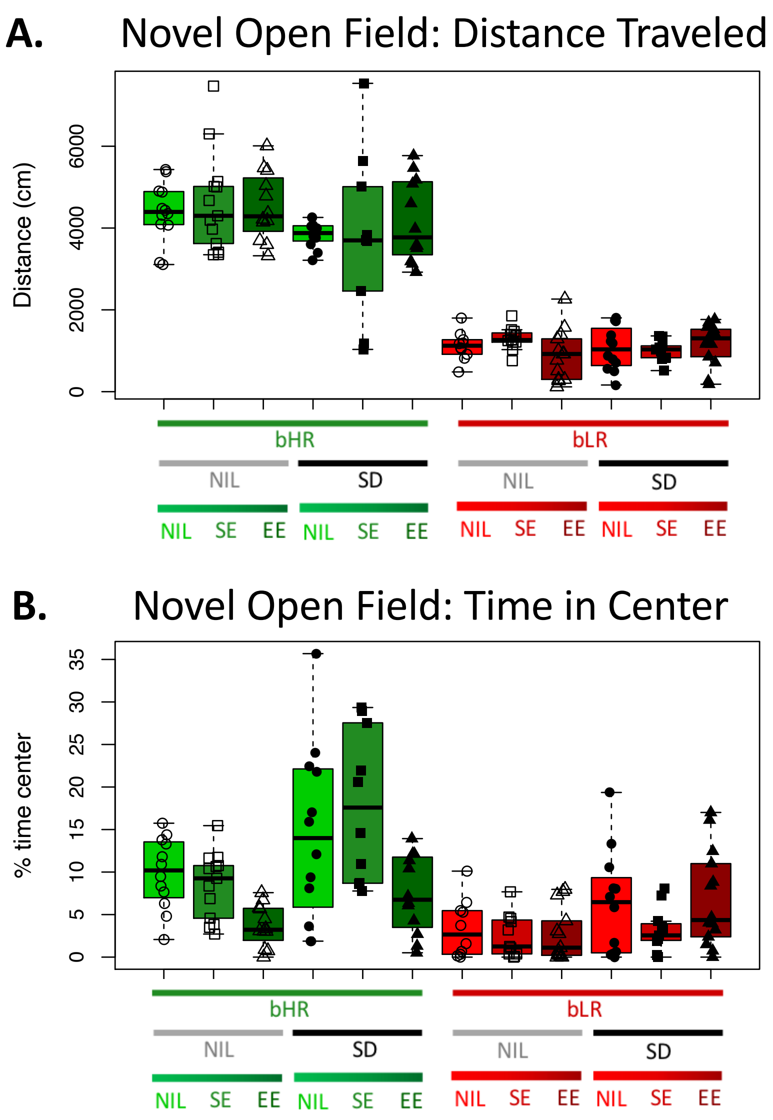

***Fig S 8. Exploratory behavior in an open field revealed expected phenotypical differences due to selective breeding, but potentially unexpected effects of social defeat and adolescent enrichment.***

*Boxplots illustrate the median and interquartile range for each treatment group (+/- whiskers illustrating the range and/or 1.5x interquartile range). Bred line is illustrated with box fill color (bHR=green, bLR=red), adolescent enrichment by datapoint shape (circle=standard housing (NIL), square=simple enrichment (SE), triangle=enhanced enrichment (EE)), and social defeat is indicated by datapoint fill (open=no defeat (NIL), black filled=defeated (SD)).* ***A.*** *bHR/bLR rats exhibited expected phenotypical differences in distance traveled in a novel open field (cm), and a small nominal decrease following social defeat* ***B.*** *bHR/bLR rats exhibited phenotypical differences in the percent time spent in the center of the open field, a nominal unexpected decrease following enrichment, and an unexpected increase following social defeat. This may be due to the animals having been placed in the center of the arena at the start of the open field habituation task, so that anxious animals that remained frozen were likely to have increased time in the center, preventing the traditional interpretation of open-field data.*

#### A panel of plasma ELISA analysis revealed little effect of behavioral interventions on testosterone, IL-6 or oxytocin

Testosterone was elevated in bLR rats in comparison to bHR rats (**Fig S9**, main effect of Line: F(1, 48)=17.5997, p=6.667e-05*), potentially more so in bLR rats kept in standard housing (interactive effects of Line*Enrichment: F(1, 48)=4.2340, p=4.480e-02#).

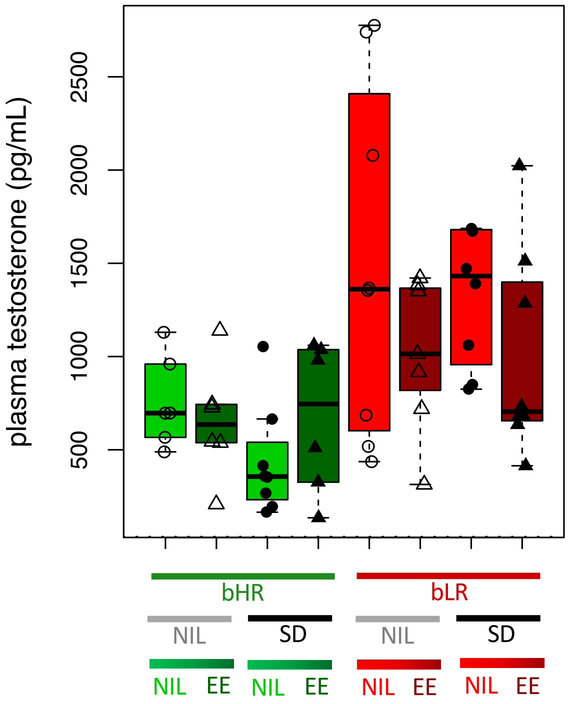

***Fig S 9. Plasma testosterone is elevated in bLR rats, potentially more so when kept in standard housing conditions.***

*Boxplot illustrates the median and interquartile range for each treatment group (+/- whiskers illustrating the range and/or 1.5x interquartile range). The dotted line represents the limit of detection for the assay. Bred line is illustrated with box fill color (bHR=green, bLR=red), adolescent enrichment by datapoint shape (circle=standard housing (NIL), triangle=enhanced enrichment (EE)), and social defeat is indicated by datapoint color (open=no defeat (NIL), black filled=defeated (SD)).*

The results from the Oxytocin and IL-6 assays were less illuminating (**Fig S10**). Neither hormone showed any effects of line, social defeat, or enrichment, but the effects may not have been fully captured by our assay due to many measurements being at or below the limit of detection (Oxytocin: 21.1%, 11/52) or near the lowest dilution on the standard curve (IL-6: 20%, 11/55, *limit of detection not provided for kit*). No conclusion could be made regarding the influence of bred line on IL-6 as the samples from the two lines were run in separate batches.

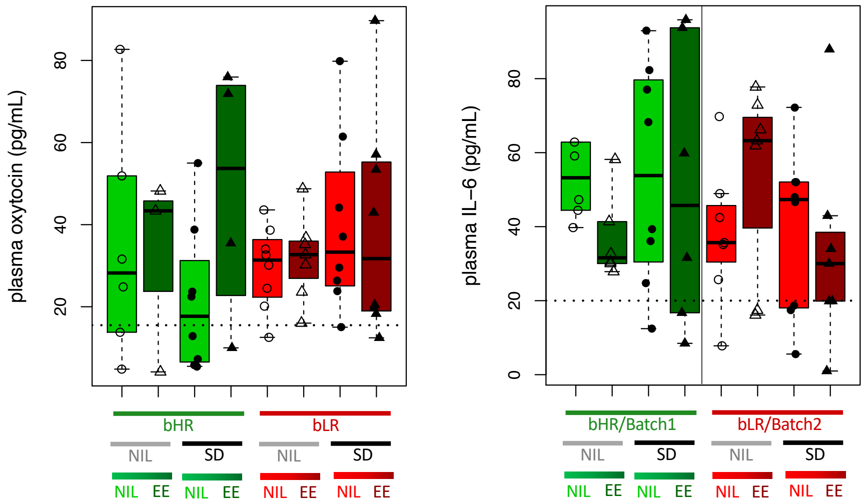

***Fig S 10. There were no significant effects of treatment group on basal plasma hormone levels of oxytocin or IL-6.***

*Boxplots illustrate the median and interquartile range for each treatment group (+/- whiskers illustrating the range and/or 1.5x interquartile range). Dotted lines represent the limit of detection for the assay, or, if this information was not available, the lowest standard dilution (IL-6). Bred line is illustrated with box fill color (bHR=green, bLR=red), adolescent enrichment by datapoint shape (circle=standard housing (NIL), triangle= enhanced enrichment (EE)), and social defeat is indicated by datapoint color (open=no defeat (NIL), black filled=defeated (SD).* ***A.*** *Plasma oxytocin (pg/mL).* ***B.*** *Plasma IL-6 (pg/mL). Note that due to technical issues, the bHR and bLR samples were run in separate batches and cannot be directly compared.*

| TOP EE GENES IN NACC (FDR<0.05) | NUCLEUS ACCUMBENS (NACC): | | | | | | | | | | | | HIPPOCAMPUS: | | | | | |
|---|---|---|---|---|---|---|---|---|---|---|---|---|---|---|---|---|---|---|
| | Ensembl_Gene_ID | Gene_Symbol | Gene_Biotype | Description | A_Log2Expression | M1_Log2FoldChange_EE | M1_Log2FoldChange_SD | M2_Log2FoldChange_EE | M2_Log2FoldChange_SD | M2_Log2FoldChange_EE*SD | Representative Enriched Gene Sets - EE | Representative Enriched Gene Sets - SD | A_Log2Expression | M1_Log2FoldChange_EE | M1_Log2FoldChange_SD | M2_Log2FoldChange_EE | M2_Log2FoldChange_SD | M2_Log2FoldChange_EE*SD |
| | ENSRNOG00000032884 | **Scn11a** | protein_coding | sodium voltage-gated channel alpha subunit 11 | 0.15 | **-2.24** | 0.44 | -1.58 | 0.84 | -1.14 | **AXON, UNUSUAL_INFECTION, ABNORMAL_VASCULAR_MORPHOLOGY** | **ABNORMAL_AGGRESSIVE_IMPULSIVE_OR_VIOLENT_BEHAVIOR, ABNORMAL_EMOTION_AFFECT_BEHAVIOR, AXON,** CATION_CHANNEL_ACTIVITY | -3.16 | 0.03 | 0.43 | -0.62 | -0.14 | 0.99 |
| | ENSRNOG00000018177 | **Dhrs2** | protein_coding | dehydrogenase/ reductase 2 | -1.70 | **-1.49** | -0.31 | **-1.63** | -0.43 | 0.32 | | | -3.62 | -0.09 | -0.17 | -0.34 | -0.40 | 0.41 |
| | ENSRNOG00000054953 | **Pcdhb6** | pseudogene | protocadherin beta 6 | 1.29 | **-1.05** | -0.11 | **-1.43** | -0.38 | 0.83 | | **NACC_NEURON** | 0.46 | -0.99 | -0.03 | -1.93 | -0.74 | 1.56 |
| | ENSRNOG00000034126 | **Pcdhga2** | protein_coding | protocadherin gamma subfamily A, 2 | 4.26 | **-0.53** | -0.08 | **-0.76** | -0.28 | 0.47 | **CELL_CELL_ADHESION, CALCIUM ION BINDING,** BONE_MARROW_STROMAL | | 4.12 | -0.54 | 0.00 | -0.89 | -0.31 | 0.58 |
| | ENSRNOG00000020084 | **Pcdhb5** | protein_coding | protocadherin beta 5 | 3.32 | **-0.31** | -0.06 | **-0.48** | -0.22 | 0.35 | **AUTISM_SPECTRUM_DISORDER_CORTEX_DOWN, SYNAPTIC_SIGNALING, OLIGODENDROCYTE_DIFFERENTIATION_DOWN, CELL_CELL_ADHESION, CALCIUM ION BINDING, MIDBRAIN_NEUROTYPES_HGABA** | **AUTISM_SPECTRUM_DISORDER_CORTEX_DOWN, SYNAPTIC_SIGNALING, OLIGODENDROCYTE_DIFFERENTIATION_DOWN, MIDBRAIN_NEUROTYPES_HGABA** & HDA2 | 3.11 | -0.42 | -0.21 | -0.73 | -0.51 | 0.53 |
| | ENSRNOG00000019276 | **Dele1** | protein_coding | DAP3 binding cell death enhancer 1 | 5.72 | -0.19 | 0.05 | **-0.34** | -0.10 | 0.30 | MITOCHONDRIAL_ENVELOPE | REGULATION_OF_HYDROLASE_ACTIVITY | 5.56 | -0.17 | -0.04 | -0.43 | -0.28 | 0.44 |
| | ENSRNOG00000046264 | **Pcdhb7** | protein_coding | protocadherin beta 7 | 3.02 | 0.23 | 0.00 | **0.40** | 0.17 | -0.37 | | **NACC_NEURON,** CA1_PYRAMIDAL_NEURON, HIPPOCAMPAL_COEXPRESSION_MOUSE_LIGHTYELLOW | 2.99 | 0.24 | 0.03 | 0.30 | 0.09 | -0.09 |
| | ENSRNOG00000020073 | **Pcdhb8** | protein_coding | protocadherin beta 8 | 3.33 | 0.45 | 0.09 | **0.73** | 0.37 | -0.56 | **CA1_CONTEXTUAL_THREAT_1 HR_POST** | **CA1_CONTEXTUAL_THREAT_1 HR_POST** | 2.46 | 0.51 | 0.22 | 1.01 | 0.71 | -0.78 |

Red=upregulated, blue=downregulated; Log2FoldChange: **Bold**=FDR<0.05, Black=p<0.05, grey=p>0.05; Gene Sets (FDR<0.05): **bold**=shared by multiple top genes

***Fig S 11. The top genes differentially expressed in relationship to EE in the NACC (FDR<0.05) often show a similar pattern of effects in the HC.***

*Results from the NACC are shown on the left, results from the HC are shown on the right. For each brain region, there are several differential expression results presented for each gene: the average log2 expression (darker grey = more highly expressed) and the log2FC associated with each targeted effect within a model only including the main effects of Enhanced Enrichment (EE) and Social Defeat (SD) ("M1") and a model including the interactive effects of EE and SD ("M2"). Blue indicates down-regulation in association with the intervention, Red indicates upregulation. Bold text indicates an effect that surpasses false discovery correction (FDR<0.05), black text indicates a nominally significant effect (p<0.05), and grey text is an effect that does not meet any threshold for significance. Representative gene sets that include the gene and were enriched for differential expression (FDR<0.05) for EE or SD in the NACC are illustrated in red (upregulated) or blue (downregulated). Bold text indicates gene sets that were either enriched for differential expression under multiple conditions (EE, SD) or included multiple top differentially expressed genes (FDR<0.05). The full differential expression results for all genes included in each of the final datasets (NACC: n=17,775, HC: n=17,629) can be found in* ***Tables S1-S2.***

| TOP SD GENES IN NACC (FDR<0.05) | NUCLEUS ACCUMBENS (NACC): | | | | | | | | | | | | HIPPOCAMPUS: | | | | | |
|---|---|---|---|---|---|---|---|---|---|---|---|---|---|---|---|---|---|---|
| | Ensembl_Gene_ID | Gene_Symbol | Gene_Biotype | Description | A_Log2Expression | M1_Log2FoldChange_EE | M1_Log2FoldChange_SD | M2_Log2FoldChange_EE | M2_Log2FoldChange_SD | M2_Log2FoldChange_EE*SD | Representative Enriched Gene Sets - EE | Representative Enriched Gene Sets - SD | A_Log2Expression | M1_Log2FoldChange_EE | M1_Log2FoldChange_SD | M2_Log2FoldChange_EE | M2_Log2FoldChange_SD | M2_Log2FoldChange_EE*SD |
| | ENSRNOG00000031065 | **AC120486.2** | unprocessed_pseudogene | RT1 class I, locus M1, 2 | -2.56 | -1.00 | -1.59 | -1.91 | **-2.67** | 2.55 | | (MHC_PROTEIN_COMPLEX) | -2.57 | -0.76 | -2.04 | -1.70 | -3.19 | 2.19 |
| | ENSRNOG00000012546 | **Frmpd1** | protein_coding | FERM and PDZ domain containing 1 | 3.44 | 0.03 | -0.22 | -0.17 | **-0.42** | 0.42 | EMBRYONIC_CORTEX_CAJAL_RETZIUS, ASTROCYTES_CORTEX_CA1, PROTEIN_LOCALIZATION_TO_MEMBRANE, NACC_NEURON_DOWN, NACC_ASTROCYTE | ASTROCYTES_CORTEX_CA1, G_PROTEIN_COUPLED_RECEPTOR_SIGNALING_PATHWAY, NACC_NEURON_DOWN, HIPPOCAMPAL_GRANULE_LAYER | 4.19 | 0.16 | -0.02 | -0.04 | -0.22 | 0.34 |
| | ENSRNOG00000039744 | **RT1-CE4** | protein_coding | RT1 class I, locus CE4 | 1.66 | -0.01 | **0.49** | 0.01 | 0.51 | -0.05 | STRIATAL_ENDOTHELIAL_CELLS_FLT1, AUTISM_SPECTRUM_DISORDER_CORTEX_UP, CELL_CELL_ADHESION, POSITIVE_REGULATION_OF_CELL_PROLIFERATION, ABNORMAL_CHOROID_MORPHOLOGY, PFC_ASTROCYTES, CHRONIC_SOCIAL_DEFEAT_UP_28D, NACC_DNMT3A_KNOCKOUT_AFTER_STRESS_DOWN, INNATE_IMMUNE_RESPONSE, EMBRYONIC_CORTEX_CAJAL_RETZIUS, ANTIGEN_PROCESSING_AND_PRESENTATION, WISTAR_KYOTO_DOWN, NACC_DNMT3A_KNOCKOUT_AFTER_STRESS_UP | STRIATAL_ENDOTHELIAL_CELLS_FLT1, RESPONSE_TO_INTERFERON_GAMMA, AUTISM_SPECTRUM_DISORDER_CORTEX_UP, PEPTIDE_ANTIGEN_BINDING, MHC_PROTEIN_COMPLEX, EMBRYONIC_CORTEX_BRAIN_B_CELL, CHRONIC_SOCIAL_DEFEAT_UP_28D, ADHD_MODEL_PCB_EXPOSURE | 2.71 | 0.14 | 0.25 | 0.63 | 0.70 | -0.76 |
| | ENSRNOG00000000512 | **Slc26a8** | protein_coding | solute carrier family 26 member 8 | 4.69 | 0.49 | **0.60** | 0.62 | 0.72 | -0.25 | CILIUM_MOVEMENT, MICROTUBULE_BASED_PROCESS | CILIUM_MOVEMENT, MICROTUBULE_BASED_PROCESS, ION_TRANSMEMBRANE_TRANSPORTER_ACTIVITY | 4.00 | 0.40 | 0.03 | 0.82 | 0.45 | -0.70 |
| | ENSRNOG00000029386 | **RT1-N2** | protein_coding | RT1 class Ib, locus N2 | 3.41 | 0.39 | **0.73** | 0.68 | **1.00** | -0.51 | | (MHC_PROTEIN_COMPLEX) | 4.56 | 0.37 | 0.56 | 1.16 | 1.30 | -1.24 |
| | ENSRNOG00000015248 | **Abca12** | protein_coding | ATP binding cassette subfamily A member 12 | 1.80 | -0.27 | 0.86 | 0.71 | **1.61** | -1.65 | UNUSUAL_INFECTION, EPITHELIUM_DEVELOPMENT, ATPASE_ACTIVITY, PROTEIN_LOCALIZATION_TO_MEMBRANE, SECRETORY_VESICLE | ABNORMAL_AGGRESSIVE_IMPULSIVE_OR_VIOLENT_BEHAVIOR, ABNORMAL_EMOTION_AFFECT_BEHAVIOR, ION_TRANSMEMBRANE_TRANSPORTER_ACTIVITY, COGNITIVE IMPAIRMENT | -0.43 | 0.17 | -0.07 | 0.08 | -0.16 | 0.14 |
| | ENSRNOG00000000723 | **RT1-CE5** | protein_coding | RT1 class I, locus CE5 | -0.96 | 0.03 | **0.96** | -0.22 | 0.75 | 0.43 | STRIATAL_ENDOTHELIAL_CELLS_FLT1, AUTISM_SPECTRUM_DISORDER_CORTEX_UP, CELL_CELL_ADHESION, POSITIVE_REGULATION_OF_CELL_PROLIFERATION, ABNORMAL_CHOROID_MORPHOLOGY, PFC_ASTROCYTES, CHRONIC_SOCIAL_DEFEAT_UP_28D, NACC_DNMT3A_KNOCKOUT_AFTER_STRESS_DOWN, INNATE_IMMUNE_RESPONSE, EMBRYONIC_CORTEX_CAJAL_RETZIUS, ANTIGEN_PROCESSING_AND_PRESENTATION, WISTAR_KYOTO_DOWN, NACC_DNMT3A_KNOCKOUT_AFTER_STRESS_UP | STRIATAL_ENDOTHELIAL_CELLS_FLT1 , RESPONSE_TO_INTERFERON_GAMMA, AUTISM_SPECTRUM_DISORDER_CORTEX_UP, PEPTIDE_ANTIGEN_BINDING, MHC_PROTEIN_COMPLEX, EMBRYONIC_CORTEX_BRAIN_B_CELL, CHRONIC_SOCIAL_DEFEAT_UP_28D, ADHD_MODEL_PCB_EXPOSURE | 0.10 | 0.17 | 0.37 | 0.50 | 0.65 | -0.48 |

Red=upregulated, blue=downregulated; Log2FoldChange: **Bold**=FDR<0.05, Black=p<0.05, grey=p>0.05; Gene Sets (FDR<0.05): **bold**=shared by multiple top genes

***Fig S 12. The top genes differentially expressed in relationship to Social Defeat (SD) in the NACC (FDR<0.05) often show a similar pattern of effects in the HC.***

*The visual table follows the same conventions as **Fig S11**. The full differential expression results for all genes included in each of the final datasets (NACC: n=17,775, HC: n=17,629) can be found in **Tables S2-S3.***

| | HIPPOCAMPUS (HC): | | | | | | | | | | | | NACC: | | | | | |
|---|---|---|---|---|---|---|---|---|---|---|---|---|---|---|---|---|---|---|
| | Ensembl_Gene_ID | Gene_Symbol | Gene_Biotype | Description | A_Log2Expression | M1_Log2FoldChange_EE | M1_Log2FoldChange_SD | M2_Log2FoldChange_EE | M2_Log2FoldChange_SD | M2_Log2FoldChange_EE*SD | Representative Enriched Gene Sets - EE | Representative Enriched Gene Sets - SD | A_Log2Expression | M1_Log2FoldChange_EE | M1_Log2FoldChange_SD | M2_Log2FoldChange_EE | M2_Log2FoldChange_SD | M2_Log2FoldChange_EE*SD |
| TOP EE GENES IN HC (FDR<0.05) | ENSRNOG00000061359 | **AABR07028009.1** | protein_coding | proteasome (prosome, macropain) activator subunit 1 (PA28 alpha), pseudo 1 | -2.47 | **-4.97** | 0.27 | **-4.71** | 0.37 | -0.44 | | | -2.48 | -2.17 | 0.02 | -2.47 | -0.16 | 0.63 |
| | ENSRNOG00000028121 | **Otop2** | protein_coding | otopetrin 2 | -1.83 | **-1.81** | -0.48 | -1.35 | -0.19 | -0.75 | | | | | | | | |
| | ENSRNOG00000000281 | **Prodh1** | protein_coding | proline dehydrogenase 1 | 5.50 | **0.48** | 0.02 | 0.35 | -0.10 | 0.22 | **PFC_ASTROCYTES, NACC_ASTROCYTE_UP, NACC_NEURON_DOWN, BRAIN_HIGH_CPG_DENSITY_PROMOTERS_WITH_H3K4ME3_AND_H3K27ME3, CHRONIC_SOCIAL_DEFEAT_UP, CA1_CONTEXTUAL_THREAT_1 HR_POST_UP, HC_CONTEXTUAL_FEAR_24HR_UP** | ABNORMALITY_OF_URINE_HOMEOSTASIS, **BRAIN_HIGH_CPG_DENSITY_PROMOTERS_WITH_H3K4ME3_AND_H3K27ME3, CHRONIC_SOCIAL_DEFEAT_UP, CA1_CONTEXTUAL_THREAT_1 HR_POST_UP,** SCHIZOPHRENIA, ACUTE_SWIM_STRESS_DOWN, **NACC_NEURON_DOWN,** HC_ASTROCYTE_GJA1, HC_COEXPRESSION_HUMAN_M8, PREFRONTAL_CORTEX_22Q11_DELETION_DOWN, HC_CONTEXTUAL_FEAR_4HR_UP | 5.04 | 0.11 | 0.01 | -0.02 | -0.13 | 0.27 |
| | ENSRNOG00000028156 | **Pld1** | protein_coding | phospholipase D1 | 4.31 | **0.49** | -0.02 | 0.29 | -0.23 | 0.35 | **EMBRYONIC_CORTEX_OLIGODENDROCYTES** | **EMBRYONIC_CORTEX_OLIGODENDROCYTES,** VACUOLE, NUCLEAR_OUTER_MEMBRANE_ENDOPLASMIC_RETICULUM_MEMBRANE_NETWORK, PARVINBETA_TARGETS_3_UP, **ABNORMAL_VASCULAR_MORPHOLOGY,** HC_COEXPRESSION_MOUSE_LIGHTCYAN, APICAL_PLASMA_MEMBRANE, LIPID_BINDING, POSTTRANSCRIPTIONAL_REGULATION_OF_GENE_EXPRESSION, PEPTIDE_METABOLIC_PROCESS, AMIDE_BIOSYNTHETIC_PROCESS, | 4.33 | 0.18 | -0.13 | 0.12 | -0.19 | 0.12 |
| | ENSRNOG00000015354 | **Aox1** | protein_coding | aldehyde oxidase 1 | 3.63 | **0.59** | -0.11 | 0.84 | 0.16 | -0.44 | **PARVINBETA_TARGETS_3_DN** | **PARVINBETA_TARGETS_3_DN** | 2.87 | 0.13 | -0.05 | 0.08 | -0.10 | 0.12 |
| | ENSRNOG00000010597 | **Slc5a7** | protein_coding | solute carrier family 5 member 7 | 1.95 | **2.06** | -1.32 | 1.84 | -1.67 | 0.44 | NIH_HIGH_ANXIETY_RATS_HIPPOCAMPUS_DOWN, **HC_CHRONIC_SOCIAL_DEFEAT,** MIDBRAIN_NEUROTYPES_HNBGABA | TRANSPORTER_ACTIVITY, **UNUSUAL_INFECTION,** EMBRYO_DEVELOPMENT, MOTOR_DELAY | 5.58 | -0.01 | 0.21 | 0.14 | 0.36 | -0.30 |

Red=upregulated, blue=downregulated; Log2FoldChange: **Bold**=FDR<0.05, Black=p<0.05, grey=p>0.05; Gene Sets (FDR<0.05): **bold**=shared by multiple top genes

***Fig S 13. The top genes differentially expressed in relationship to Enhanced Enrichment (EE) in the HC (FDR<0.05) often show a similar pattern of effects in the NACC.***

*Results from the HC are shown on the left, results from the NACC are shown on the right. Otherwise, the visual table follows the same conventions as* ***Fig S11****. The full differential expression results for all genes included in each of the final datasets (NACC: n=17,775, HC: n=17,629) can be found in* ***Tables S2-S3.***

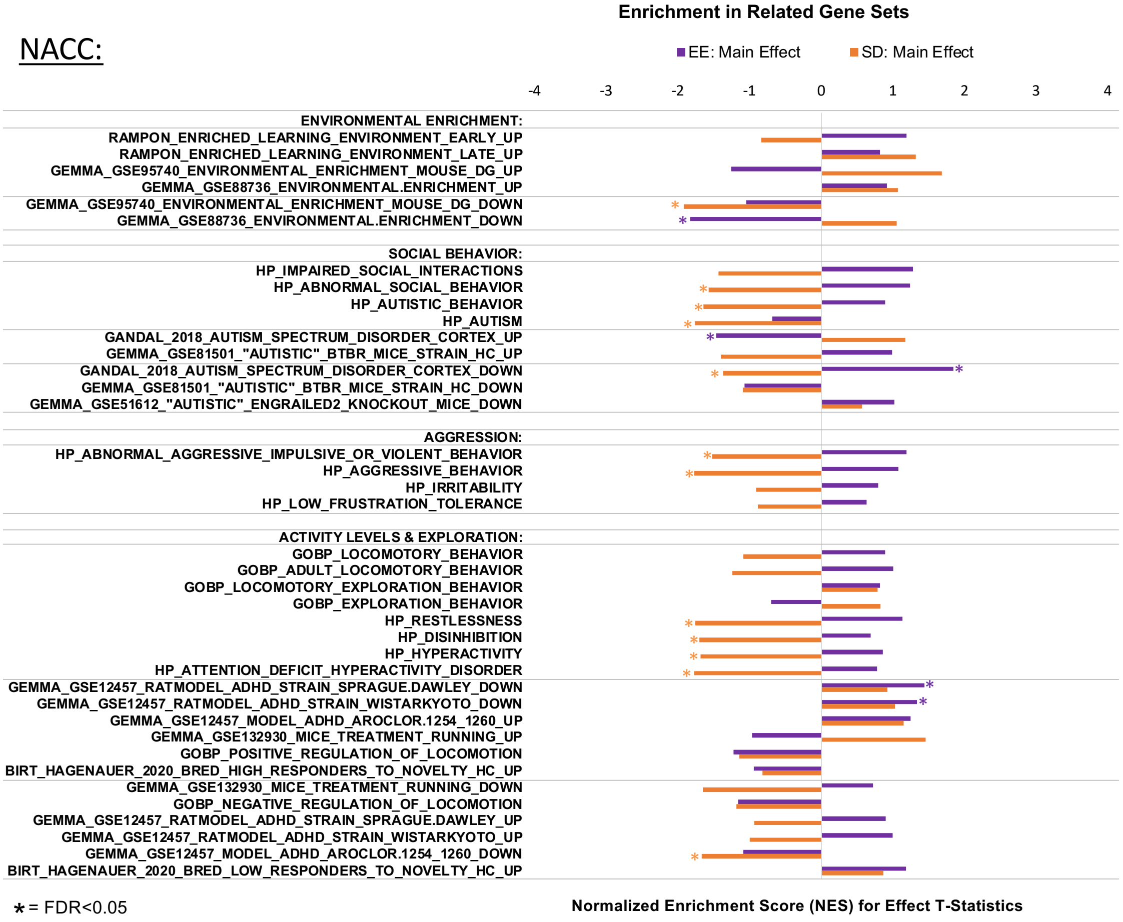

***Fig S 14. Gene sets that were previously associated with social behavior, aggression, and locomotor activity are enriched with down-regulated expression related to Social Defeat (SD) in the NACC.***

*Gene set enrichment analysis was run using the t-statistics for each condition (Enhanced Enrichment (EE), SD) outputted from the model targeting main effects ("M1") and a custom gene set file (.GMT). Bars indicate the Normalized Enrichment Score (NES) for each gene set, with positive indicating enrichment with upregulation and negative indicating enrichment for downregulation. Orange indicates the NES for SD effects, purple for EE effects. Asteriskes indicate that the enrichment within a gene set has surpassed false discovery correction (FDR<0.05). Grey lines are used to visually denote gene sets within each broad category of behavior by the direction of their previously-defined association with that behavior: directionless, gene sets previously associated with increases in that behavior, and gene sets previously associated with decreases in that behavior.*

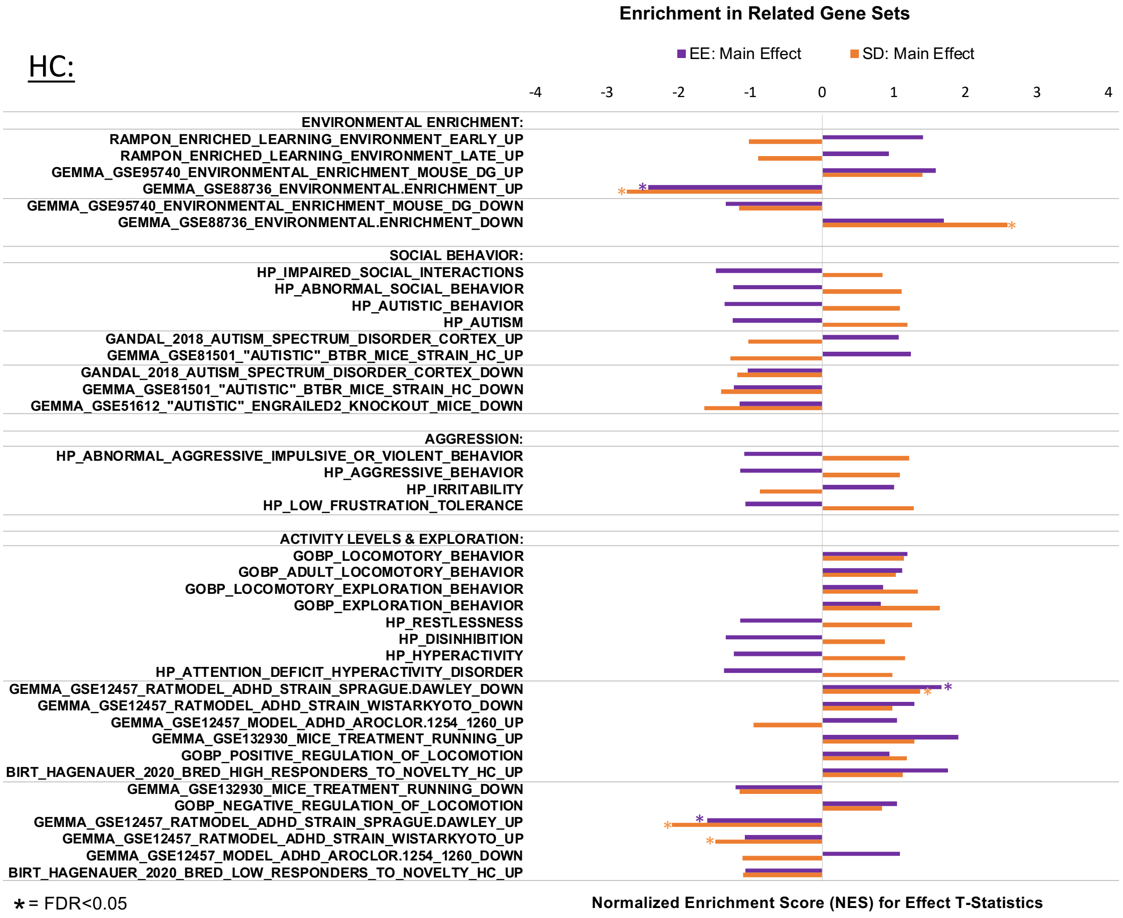

***Fig S 15. Gene sets that were previously associated with increased locomotor activity are enriched with upregulation related to Enhanced Enrichment (EE) and Social Defeat (SD) in the HC, and gene sets associated with decreased locomotor activity are enriched with down-regulation.***

*Plotting conventions follow those of **Fig S14**.*

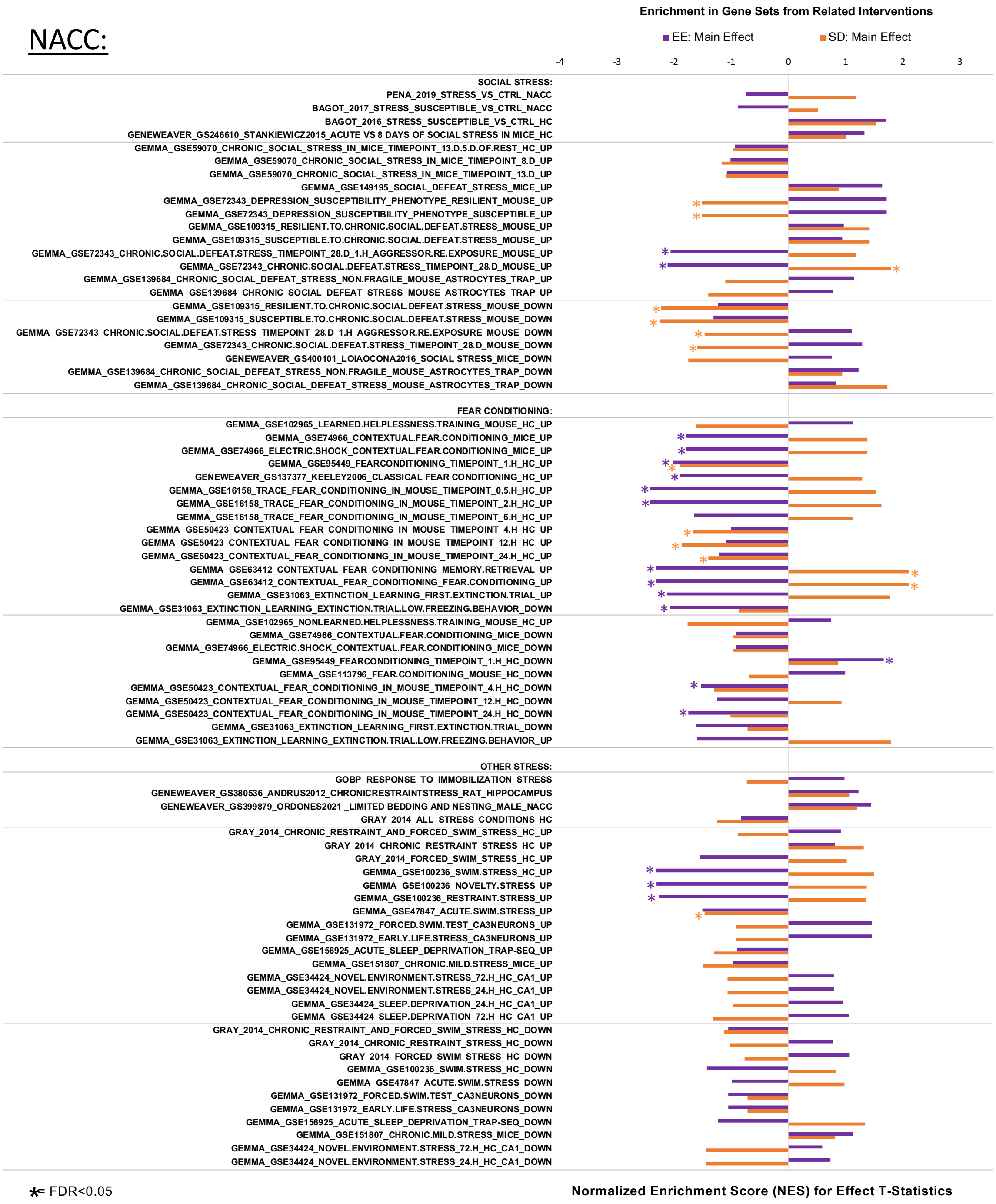

***Fig S 16. Gene sets that were previously associated with social stress, fear conditioning, and other stress are highly enriched with differential expression related to Enhanced Enrichment (EE) and Social Defeat (SD) in the NACC.***

*The associations with EE and SD in these gene sets are often in opposing directions. Plotting conventions follow those of **Fig S14.***

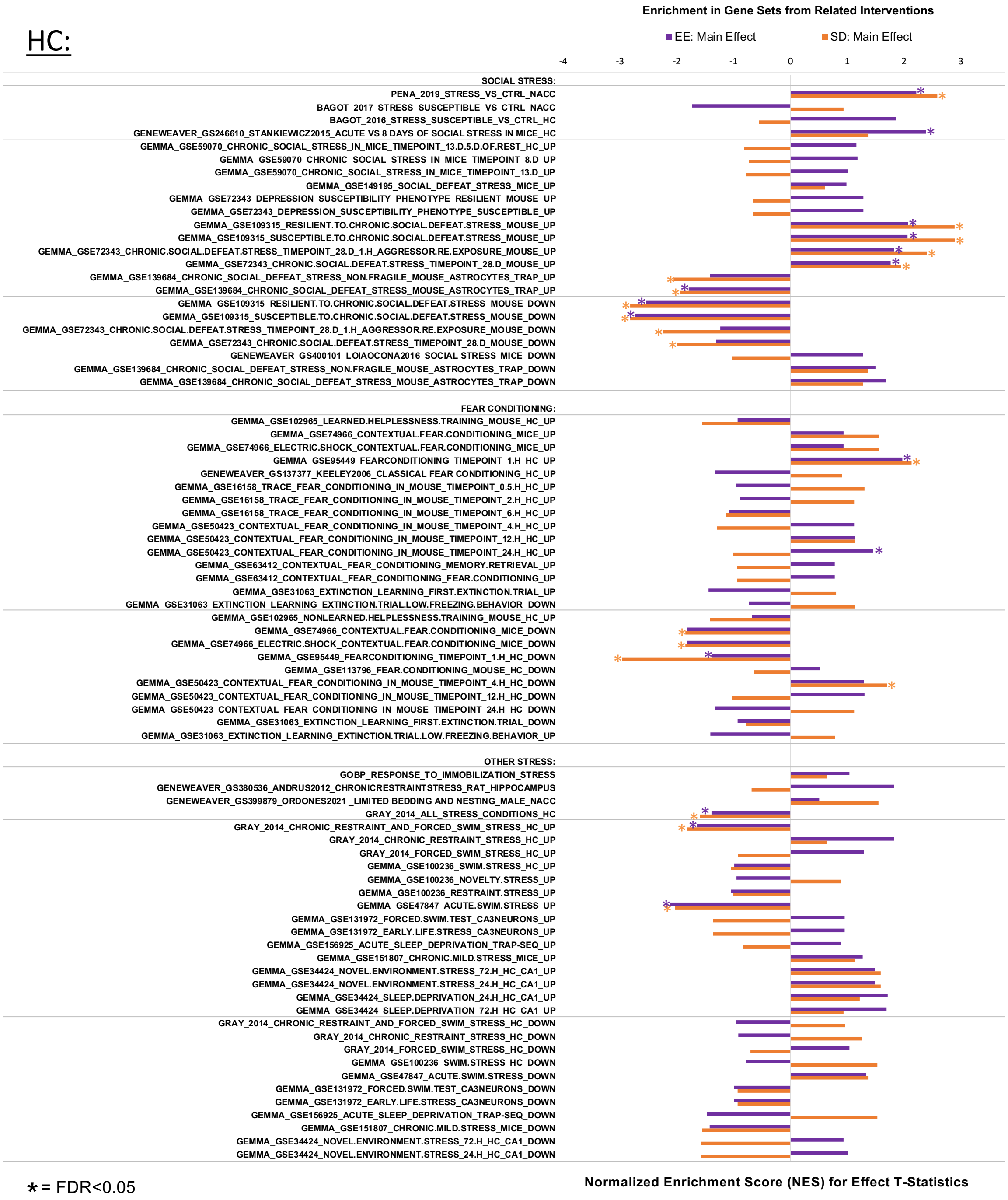

***Fig S 17. Gene sets that were previously associated with social stress, fear conditioning, and other stress are highly enriched with differential expression related to Enhanced Enrichment (EE) and Social Defeat (SD) in the HC.***

*The associations with EE and SD in these gene sets are typically in the same direction. Plotting conventions follow those of* ***Fig S14****.*

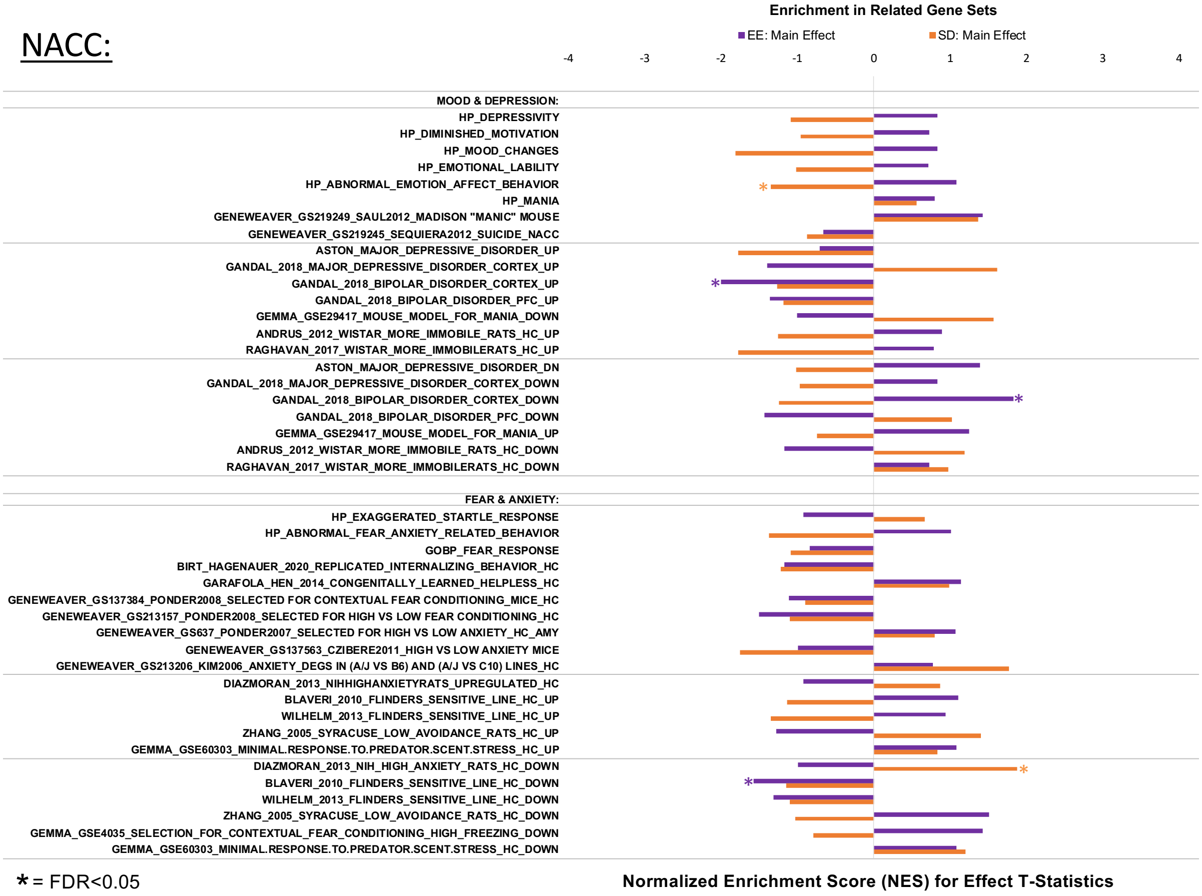

***Fig S 18. A few gene sets related to mood/depression and endogenous fear & anxiety were enriched for differential expression related to Social Defeat (SD) and Enhanced Enrichment (EE) in the NACC.***

*Plotting conventions follow those of* ***Fig S14.***

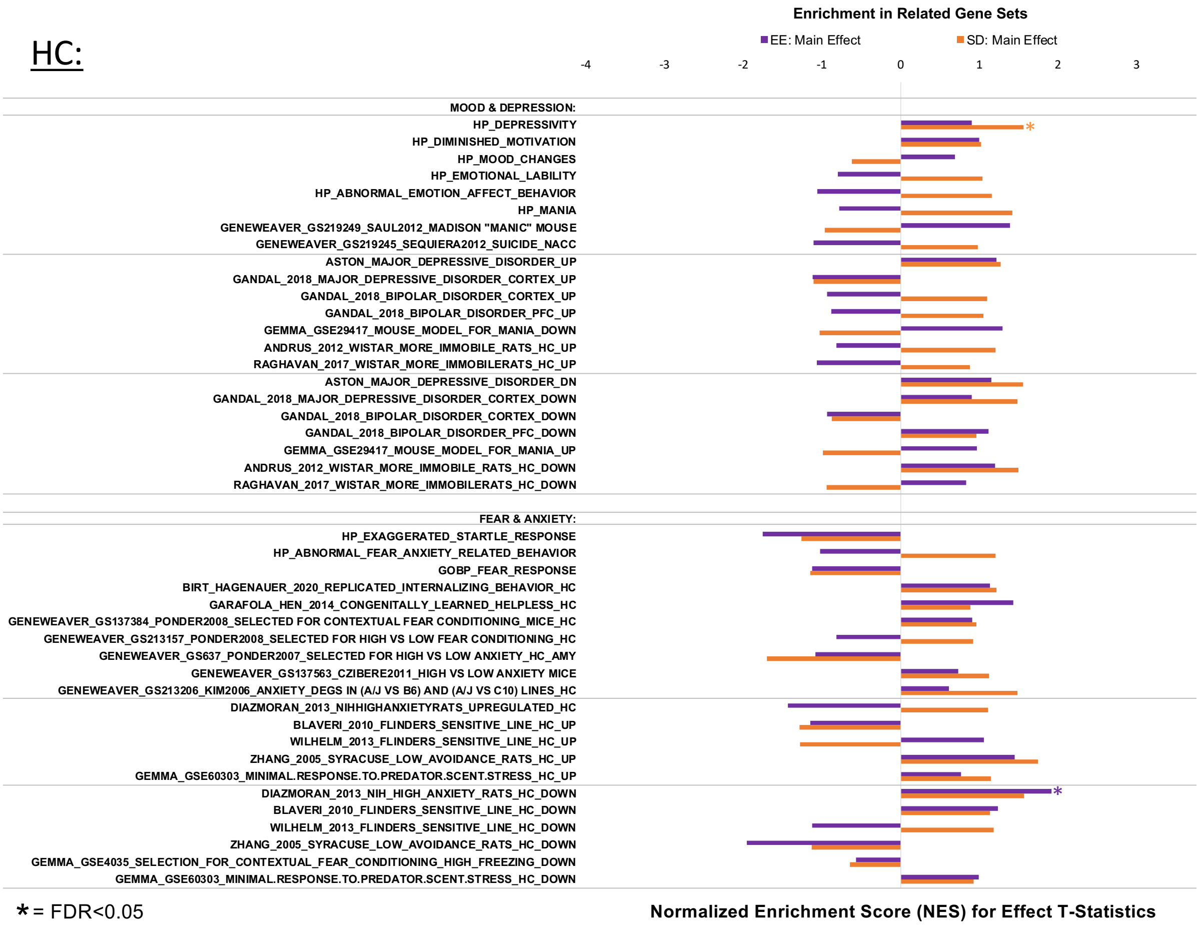

***Fig S 19. A few gene sets related to mood/depression and endogenous fear & anxiety were enriched for differential expression related to Social Defeat (SD) and Enhanced Enrichment (EE) in the HC.***

*Plotting conventions follow those of* ***Fig S14.***

NACC:

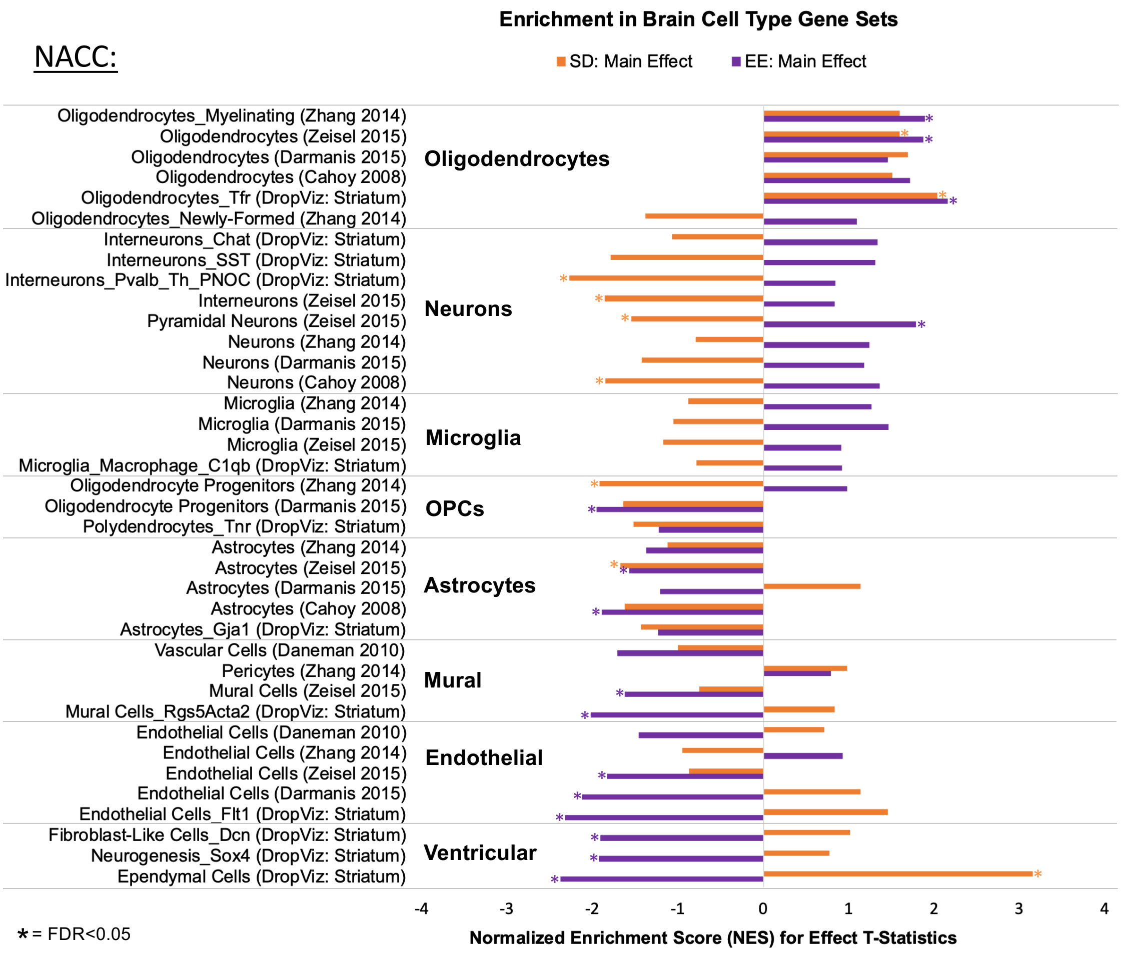

***Fig S 20. Cell type related gene sets are enriched with differential expression in the NACC associated with SD and EE.***

*Gene sets related to oligodendrocytes were enriched with upregulation in the NACC in association with both SD and EE. Gene sets related to neurons were enriched with down-regulation in association with SD**,** and gene sets related to a variety of cell types associated with the brain vascular and ventricular systems (astrocytes, mural cells, endothelial cells, oligodendrocyte progenitor cells (OPCs) and neurogenesis-related cells, ependymal cells) were enriched with down-regulation associated with EE. Enrichment within other cell type-related gene sets were less consistent across gene sets representing the same category. Plotting conventions follow those of **Fig S14.***

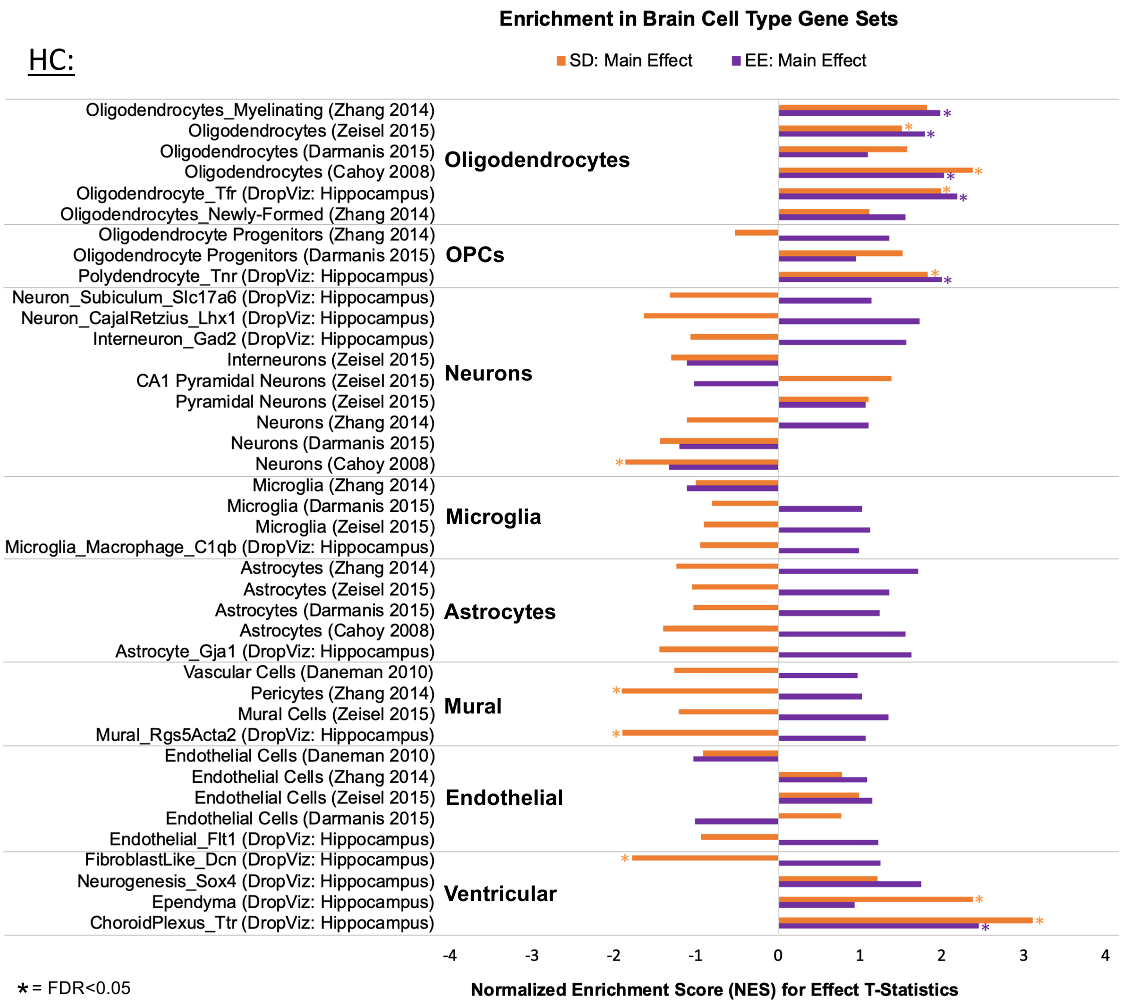

***Fig S 21. Cell type related gene sets are enriched with differential expression in the HC associated with SD and EE.***

*Similar to the NACC, gene sets related to oligodendrocytes were enriched with upregulation in the HC in association with both SD and EE. Two gene sets related to mural cells were enriched with down-regulation in the HC following SD, and gene sets related to the choroid plexus were upregulated in the HC in association with both conditions. Enrichment within other cell type-related gene sets were less consistent across gene sets representing the same category. Plotting conventions follow those of* ***Fig S14.***

| | Gene Symbol | A_Log2Expression | M1_Log2FoldChange_EE HC | M1_Log2FoldChange_SD HC | bLR vs. bHR: Cohen's D (Birt et al. 2021) |
|---|---|---|---|---|---|
| EE in HC FDR<0.05 | AABR07028009.1 | -2.47 | **-4.97** | 0.27 | NA |
| | Otop2 | -1.83 | **-1.81** | -0.48 | NA |
| | Prodh1 | 5.50 | **0.48** | 0.02 | NA |
| | Pld1 | 4.31 | **0.49** | -0.02 | -0.48 |
| | Aox1 | 3.63 | **0.59** | -0.11 | 0.14 |
| | **Slc5a7** | 1.95 | **2.06** | -1.32 | -0.76 |
| EE in NACC FDR<0.05 | Scn11a | -3.16 | 0.03 | 0.43 | *-0.57* |
| | **Dhrs2** | -3.62 | -0.09 | -0.17 | 1.20 |
| | Pcdhb6 | 0.46 | -0.99 | -0.03 | 0.43 |
| | Pcdhga2 | 4.12 | -0.54 | 0.00 | 0.22 |
| | **Pcdhb5** | 3.11 | -0.42 | -0.21 | 1.12 |
| | Dele1 | 5.56 | -0.17 | -0.04 | *-0.74* |
| | Pcdhb7 | 2.99 | 0.24 | 0.03 | -0.57 |
| | Pcdhb8 | 2.46 | 0.51 | 0.22 | *-0.77* |
| SD in NACC FDR<0.05 | AC120486.2 | -2.57 | -0.76 | -2.04 | 0.55 |
| | **Frmpd1** | 4.19 | 0.16 | -0.02 | -0.89 |
| | RT1-CE4 | 2.71 | 0.14 | 0.25 | 0.41 |
| | Slc26a8 | 4.00 | 0.40 | 0.03 | -0.49 |
| | RT1-N2 | 4.56 | 0.37 | 0.56 | -0.51 |
| | Abca12 | -0.43 | 0.17 | -0.07 | 0.01 |
| | RT1-CE5 | 0.10 | 0.17 | 0.37 | -0.35 |

***Fig S 22. Some of the top differentially expressed genes in our study have been previously shown to have nominal (p<0.05) bLR/bHR differential expression in the hippocampus.***

*The table includes the top DEGs (FDR<0.05) from our study for the effects of adolescent enrichment (EE) and social defeat (SD) in the nucleus accumbens (NACC) and hippocampus (HC). For reference, for each of the DEGs the first three columns include the Log2 expression levels in the HC and the Log2FoldChange for the main effects of EE and SD in the HC within our study (using the main effects model: M1), copied from* ***Figs S12-14****. Positive values (red) indicate upregulation with the condition, negative values (blue) indicated down-regulation. The final column shows the estimated Cohen's d for the bLR vs. bHR differential expression in the HC from a published cross-generational meta-analysis [18] Positive values (red) indicate elevated expression in bLRs (vs. bHRs), negative values (green) indicate elevated expression in bHRs (vs. bLRs). For all columns, grey text indicates no significant effect in the HC (p>0.10),* *black italicized text indicates a statistical trend in the HC (0.05<p<0.10), black text indicates a nominally significant effect in the HC (p<0.05), and black bold text indicates an effect surviving false discovery rate correction in the HC (FDR<0.05). Note that more than half of the DEGs for EE that were included in the bHR/bLR meta-analysis (6 out of 11) show indications of possible bLR/bHR differential expression (p<0.10).*

### Supplementary Table Legends

##### *Table S 1. An excel file (.xlsx) containing the detailed statistical reporting for the behavioral tests, hormonal assays, and gene set category enrichment results.*

The file contains four spreadsheets containing the detailed statistical reporting for four types of dependent variables: 1) Social Defeat Behavior, 2) Behavioral Battery, 3) Hormonal Assays, 4) Enrichment in Gene Set Categories, as well as a fifth spreadsheet containing a description of how litters were distributed amongst the treatment groups. A description of the statistical test used for each type of dependent variable is at the top of each of the statistical reporting spreadsheets. Each dependent variable is also labeled with the Figure number where the results are visualized in the manuscript or supplement. *Abbreviations:* Chisq=Chi-Square Statistic, Df=Degrees of Freedom, p=p-value, adj.p=p-value adjusted for multiple comparisons using the Bonferonni method, #=nominally significant (p<0.05), **=significant (adj.p<0.05), SS=Sum of Squares, F=F-statistic, Variable:Variable=an effect due to the interaction of two independent variables, Variable:Variable:Variable=an effect due to the interaction of three independent variables, OR=Odds Ratio, NACC=Nucleus Accumbens, HC=Hippocampus, EE= Enhanced Enrichment, SD=Social Defeat.

##### *Table S 2. An excel file (.xlsx) containing the full differential expression results for the NACC (17,775 Ensembl-annotated genes).*

The first spreadsheet provides the output for a model containing only the main effects of Enhanced Enrichment and Social Defeat along with technical co-variates ("M1"), the second spreadsheet provides the output for a model containing the interactive effects of Enhanced Enrichment and Social Defeat along with technical co-variates ("M2"). For each row in the spreadsheets, annotation is provided in the form of Ensembl gene ID and official gene symbol. The statistical output for each gene includes average log2 cpm expression ("A"), the log2FC ("Coef"), the t-statistic ("t."), the nominal p-value ("p.value"), and the false discovery rate (FDR) corrected p-value ("p.value.adj") for each of the variables of interest. The rows are ordered by Ensembl ID. Conditional formatting is used to make the direction of effect easy to see, with pink indicating upregulation (a positive Log2FC Coef) and blue indicating downregulation (a negative Log2FC Coef). Conditional formatting is also used to make significant effects easier to see, with green indicating a low nominal p-value or low FDR (adj.p-value), and red indicating a high nominal p-value or high FDR.

***Table S 3. An excel file (.xlsx) containing the full differential expression results for the HC (17,629 Ensembl-annotated genes).***

The first spreadsheet provides the output for a model containing only the main effects of Enhanced Enrichment and Social Defeat along with technical co-variates ("M1"), the second spreadsheet provides the output for a model containing the interactive effects of Enhanced Enrichment and Social Defeat along with technical co-variates ("M2"). For each row in the spreadsheets, annotation is provided in the form of Ensembl gene ID and official gene symbol. The statistical output for each gene includes average log2 cpm expression ("A"), the log2FC ("Coef"), the t-statistic ("t."), the nominal p-value ("p.value"), and the false discovery rate (FDR) corrected p-value ("p.value.adj") for each of the variables of interest. The rows are ordered by Ensembl ID. Conditional formatting is used to make the direction of effect easy to see, with pink indicating upregulation (a positive Log2FC Coef) and blue indicating downregulation (a negative Log2FC Coef). Conditional formatting is also used to make significant effects easier to see, with green indicating a low nominal p-value or low FDR (adj.p-value), and red indicating a high nominal p-value or high FDR.

***Table S 4. An excel file (.xlsx) containing the full gene set enrichment (fGSEA) results for all gene sets included in the final NACC analysis (n=10,484 gene sets).***

The first spreadsheet ("NAcc_All_fGSEA_Results") contains the results for the gene set enrichment (fGSEA) analysis of the effects of adolescent Enhanced Enrichment and Social Defeat as calculated by either preranking the genes using using the differential expression results (t-statistics) from the main effects model (M1) or the Interactive Effects Model (M2). For each row, the gene set name is given in the "pathway" column. The variable (Enhanced Enrichment (EE) or Social Defeat (SD)) and model (M1 or M2) are indicated in row 1 for each set of fGSEA statistics. Each set of fGSEA results includes a nominal p-value ("pval"), false discovery rate (FDR) corrected p-value ("padj"), enrichment score ("ES"), normalized enrichment score ("NES"), number of genes in the gene set with effects beyond what would be expected by random chance (as determined by permutation, "nMoreExtreme"), gene set size, and a list of the genes in the gene set with the largest effects ("leading genes"). Conditional formatting is used to make the direction of effect easy to see, with pink indicating an enrichment of upregulation in the gene set (a positive ES and NES) and blue indicating an enrichment of downregulation (a negative ES and NES). Conditional formatting is also used to make significant effects easier to see, with green indicating a low nominal p-value or low FDR (adj.p-value), and red indicating a high nominal p-value or high FDR. The final columns contain the minimum adjusted p-value for both sets of results (from M1 and M2) for the variable of Enrichment ("EE_Min_AdjPval") and the variable of Social Defeat ("SD_MIN_AdjPval"). The values in these columns are used to provide two additional worksheets that are sorted to illustrate the top enriched gene sets for the variables of Enrichment ("SortByEE") and Social Defeat ("SortBySD").

***Table S 5. An excel file (.xlsx) containing the full gene set enrichment (fGSEA) results for all gene sets included in the final HC analysis (n=10,540 gene sets).***

The first spreadsheet (“HC_All_fGSEA_Results”) contains the results for the gene set enrichment (fGSEA) analysis of the effects of adolescent Ehanced Enrichment and Social Defeat as calculated by either preranking the genes using using the differential expression results (t-statistics) from the main effects model (M1) or the Interactive Effects Model (M2). For each row, the gene set name is given in the “pathway” column. The variable (Enhanced Enrichment (EE) or Social Defeat (SD)) and model (M1 or M2) are indicated in row 1 for each set of fGSEA statistics. Each set of fGSEA results includes a nominal p-value (“pval”), false discovery rate (FDR) corrected p-value (“padj”), enrichment score (“ES”), normalized enrichment score (“NES”), number of genes in the gene set with effects beyond what would be expected by random chance (as determined by permutation, “nMoreExtreme”), gene set size, and a list of the genes in the gene set with the largest effects (“leading genes”). Conditional formatting is used to make the direction of effect easy to see, with pink indicating an enrichment of upregulation in the gene set (a positive ES and NES) and blue indicating an enrichment of downregulation (a negative ES and NES). Conditional formatting is also used to make significant effects easier to see, with green indicating a low nominal p-value or low FDR (adj.p-value), and red indicating a high nominal p-value or high FDR. The final columns contain the minimum adjusted p-value for both sets of results (from M1 and M2) for the variable of Enrichment (“EE_Min_AdjPval”) and the variable of Social Defeat (“SD_MIN_AdjPval”). The values in these columns are used to provide two additional worksheets that are sorted to illustrate the top enriched gene sets for the variables of Enrichment (“SortByEE”) and Social Defeat (“SortBySD”).
